# Human α-Synuclein Inhibits Platelets Aggregation *in vitro* by Interfering with the α-Thrombin/Protease-Activated Receptor 1 Functional Axis

**DOI:** 10.1101/2021.03.28.437436

**Authors:** Giulia Pontarollo, Laura Acquasaliente, Claudia Maria Radu, Daniele Peterle, Ilaria Artusi, Anna Pagotto, Federico Uliana, Paolo Simioni, Alessandro Negro, Vincenzo De Filippis

## Abstract

α-Synuclein (αSyn) is a small (140 amino acids) disordered, acidic (pI: 4.7) protein, highly conserved in vertebrates and implicated in the pathogenesis of Parkinson’s disease (PD), a neurodegenerative disease characterized by the deposition of αSyn amyloid fibrils in dopaminergic neurons. Beyond the central nervous system, significant expression of αSyn has also been measured in the blood (~1 μM), where platelets are the main cellular hosts of αSyn. Although the pathological implication of αSyn in PD is widely accepted, the physiological role of blood αSyn is still elusive. Starting from the notion that platelets are either the major cellular reservoir of αSyn in the blood and, concomitantly, act as key players in hemostasis, being activated also by α-thrombin (αT) *via* cleavage of protease-activated receptors (PARs), we decided to investigate the possibility that αSyn could modulate platelet activation by interfering with the αT-PAR functional axis. Using multiple electrode aggregometry, i.e. a fast and specific platelet-function-testing method, as well as steady-state fluorescence spectroscopy, surface plasmon resonance, and fluorescence microscopy, we show here that monomeric αSyn functions as a negative regulator of αT-mediated platelets activation. αSyn acts either directly, *via* competitive inhibition of PAR1 activation by αT and TRAP6 agonist, and indirectly, by scavenging αT on the platelet plasma membrane. A simple electrostatic model of αSyn platelet antiaggregating effect is proposed and the possible role of the protein at the interplay of amyloidosis and thrombosis is discussed.

---

α-Synuclein (αSyn) is a small (140 amino acids; ~14 kDa) acidic protein, highly conserved in vertebrates (1) and implicated in the pathogenesis of Parkinson’s disease (PD) (2). αSyn is a structurally disordered monomeric protein, either when isolated in solution (3) or in cellular environments, where it assumes a loosely packed, dynamic structure (4). αSyn primary structure displays three distinctive regions (**Fig. 1**): i) the N-terminal region (NT, amino acids 1-60) is highly electropositive, and serves to preferentially localize αSyn onto negatively charged biological membranes (5); ii) the central region, corresponding to the Non-Amyloid β-Component (NAC, amino acids 61-95) is hydrophobic in nature and crucial for fibrillation (6); iii) the C-terminal region (CT, amino acids 96-140) displays high electronegative potential and is responsible of αSyn binding to several target proteins (7). Upon prolonged incubation, αSyn aggregates and forms amyloid fibrils (8), characterized by a cross-β-sheet structure and stabilized by extensive hydrogen bonds network (9). The kinetics of αSyn amyloid formation follows a nucleation-elongation mechanism (3) with a critical αSyn concentration of 28 μM after 3-day incubation at 37°C in TBS, pH 7.5 (8). The lag phase of fibril formation is significantly shortened at higher αSyn concentrations (10) and in crowded environments (11).

**Figure 1.**
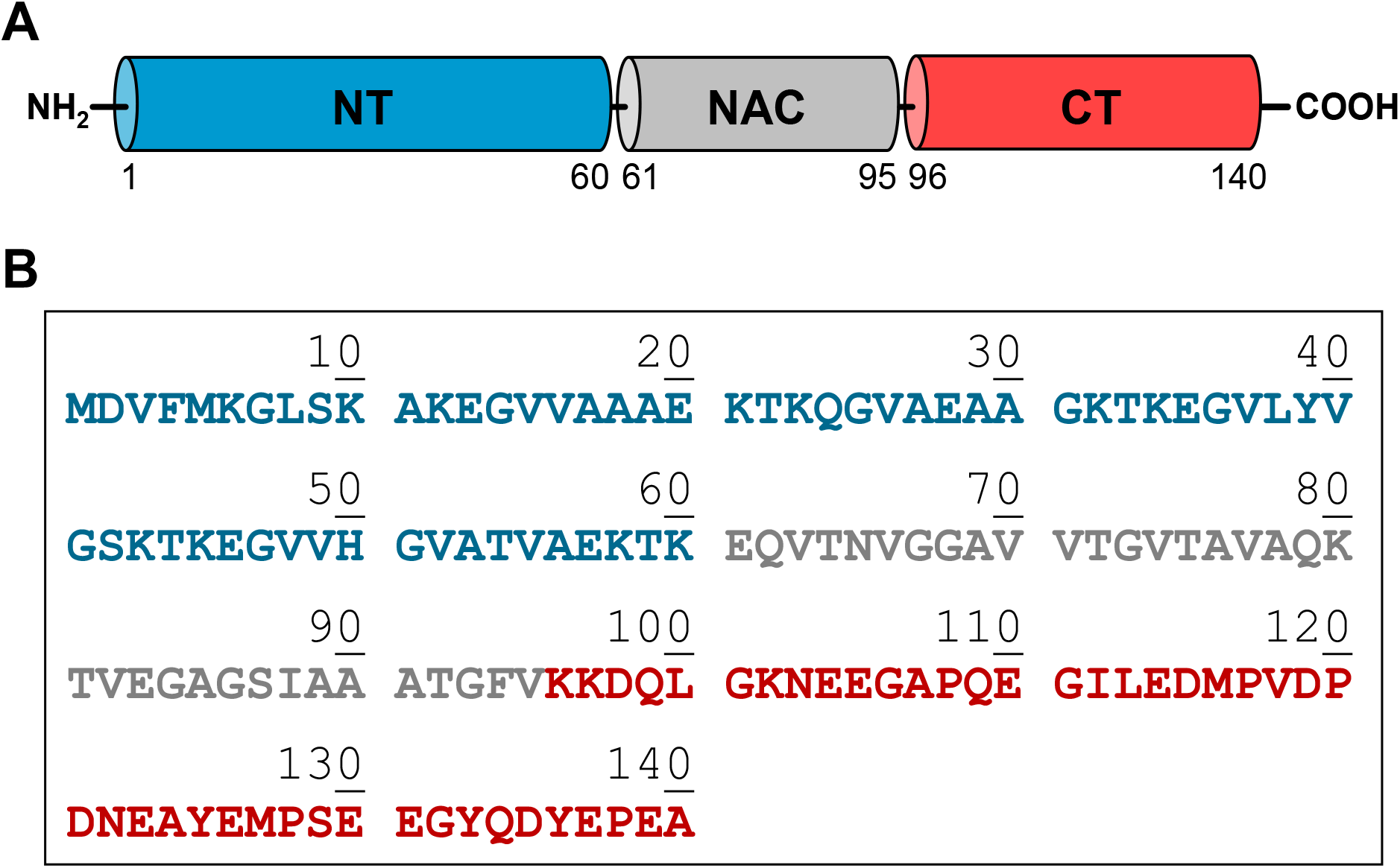
Domain architecture (A) and amino acid sequence (B) of human αSyn. NT: the N-Terminal domain region (amino acids 1-60) is positively charged (pI: 9.4) and assumes a helical conformation on lipid membrane surfaces. NAC: the Non-Amyloid β-Component (amino acids 61-95) is highly hydrophobic, has strong β-sheet conformational propensity, and mediates αSyn aggregation/fibrillation. CT: the C-Terminal Domain (amino acids 96-140), is strongly negative (pI: 3.1).

Clinical evidences indicate that αSyn plays a major role in the pathogenesis of Parkinson’s disease (PD) and the presence of αSyn amyloid aggregates (i.e. Lewy bodies) in the dopaminergic neurons of the brain *substantia nigra* is a key neuropathological hallmark of PD (2,12,13). Whereas the role of αSyn in the pathogenesis of PD is widely accepted (14,15), the physiological role of this protein is still elusive (16). αSyn is abundantly present *in vivo* in the human central nervous system, in the nuclei of neuronal cells and presynaptic terminals, where it binds to synaptic vesicles and modulate vesicle homeostasis and synaptic plasticity (17). Beyond central nervous system, significant expression levels of αSyn have also been measured in the cerebrospinal fluid (18,19), haematopoietic tissue (20,21) and blood (~1 μM; ~15 mg/l) (22–25). The vast majority (>99%) of blood αSyn is found in the erythrocytes (as absolute amount), while the remaining is split between plasma (0.1%), leukocytes (0.05%) and platelets (0.2%), where the latter are the main cellular hosts of αSyn in the blood (23,26). Although αSyn has been abundantly found in the cytoplasm of resting platelets, associated with the membrane of secretory α-granules and the inner leaflet of the plasma membrane (20,23,24), little is known about the (extracellular) localization of the protein after platelets activation. Notably, exogenous αSyn has been shown to pass across platelet plasma membrane and impair αT-stimulated α-granule release (27).

Platelets play a pivotal role in primary haemostasis, as they adhere to and are activated by subendothelial matrix proteins (i.e. collagen and von Willebrand factor) that become exposed after vascular injury, thus forming the primary platelet plug (28). Activated platelets then undergo dramatic shape change, associated with dense and α-granules secretion and inversion of plasma membrane polarity, accomplished *via* phosphatidylserine exposure, which is instrumental for coagulation factors complex assembly and activation (i.e. the tenase and prothrombinase complexes) during the amplification and propagation phases of αT generation (28).

Noteworthy, thrombin (αT) is the most potent activator of platelets *in vivo*, cleaving the exodomain of two G-protein-coupled receptors (GPCRs) on the platelets surface, i.e. type-1 and type-4 Protease Activated Receptors (PARs) (29). After cleavage, the newly generated N-termini act as intramolecular activators of PARs, finally leading to degranulation and morphological and functional changes typical of activated platelets (29). At variance with αT, ADP activates platelets by directly interacting with P2Y_12_ receptor (i.e. the major GPCR for ADP on platelets membrane), reducing cAMP concentration and increasing cytosolic Ca^2+^, with final platelets shape change and activation (30).

Starting from the notion that platelets are either the main cellular reservoir of αSyn in the blood and, concomitantly, are the main cellular target of αT procoagulant activity, we decided to investigate the possibility that αSyn could modulate platelet activation by interfering with the αT-PAR functional axis. Our results concurrently indicate that αSyn functions as a negative regulator of αT-mediated platelets activation, acting either directly, *via* competitive interaction with PAR1 in αT binding, and indirectly, by scavenging αT on platelets plasma membrane. The possible physio-pathological implications of these findings will be also discussed.

## Results

### Production and characterization of recombinant αSyn species

Human full-length α-synuclein (αSyn, amino acids 1-140), the corresponding N-terminal 6xHis-tag derivative (6xHis-αSyn), the truncated species 6xHis-αSyn(1-96) and the fusion mutant protein αSyn-GFP, in which the polypeptide chain sequence of the Green-Fluorescent Protein (GFP) was fused with αSyn C-terminal end, were expressed in *Escherichia coli* cells, as previously detailed (31,32). For wild-type αSyn and αSyn-GFP protein mutant, the bacterial pellet was sonicated and then boiled for 10 min. After centrifugation, the supernatant enriched with αSyn or αSyn-GFP was dialyzed and further purified by RP-HPLC. At variance, 6xHis-αSyn and 6xHis-αSyn(103-140) were purified by immobilized metal ion affinity chromatography (IMAC), followed by RP-HPLC. The purity of αSyn species was checked by SDS-PAGE and RP-HPLC (>98%), while their chemical identity established by high-resolution mass spectrometry, which was found in agreement with the protein amino acid composition within 20 ppm mass accuracy (**Supplementary Table S1 and Figure S1**). Monomerization of purified αSyn was achieved by alkaline treatment, i.e. dissolution of αSyn lyophilizate with NaOH solution, at pH 11.0, followed by addition of 0.1 M Tris-HCl, pH 7.0, down to pH 8.0 (33). The monomeric state of αSyn preparation was confirmed by dynamic light scattering measurements (DLS), from which a hydrodynamic diameter (d_H_) of 56 ± 6 Å was estimated, with a percent polydispersity (%PD) as low as 11.6% (**Supplementary Figure S1**), indicative of a monodispersed protein solution. Of note, the size of αSyn reported in this study is lower than that predicted for a fully unfolded protein of 140 amino acids like αSyn (d_HU_ = 68 Å), but nevertheless it compares favourably with that determined experimentally by small-angle X-ray scattering (d_H_ = 54 ± 2 Å) and size-exclusion chromatography (d_H_ = 55 ± 6 Å) (33), and is in keeping with the loosely packed, dynamic structure recently elucidated for monomeric αSyn (4).

### Inhibition of platelet aggregation by αSyn and its derivatives

Multiple Electrode Aggregometry (MEA) was used to estimate the effect of full-length αSyn and its N- and C-terminally truncated species on platelets aggregation induced in whole blood or isolated platelets by TRAP6, αT or ADP (**Fig. 2**). Notably, TRAP6 is the synthetic hexapeptide SFLLRN-NH_2_, corresponding to the N-terminal segment of the tethered PAR1 region, acting as a potent and specific PAR1 agonist independently of αT-induced proteolysis (34).

**Figure 2.**
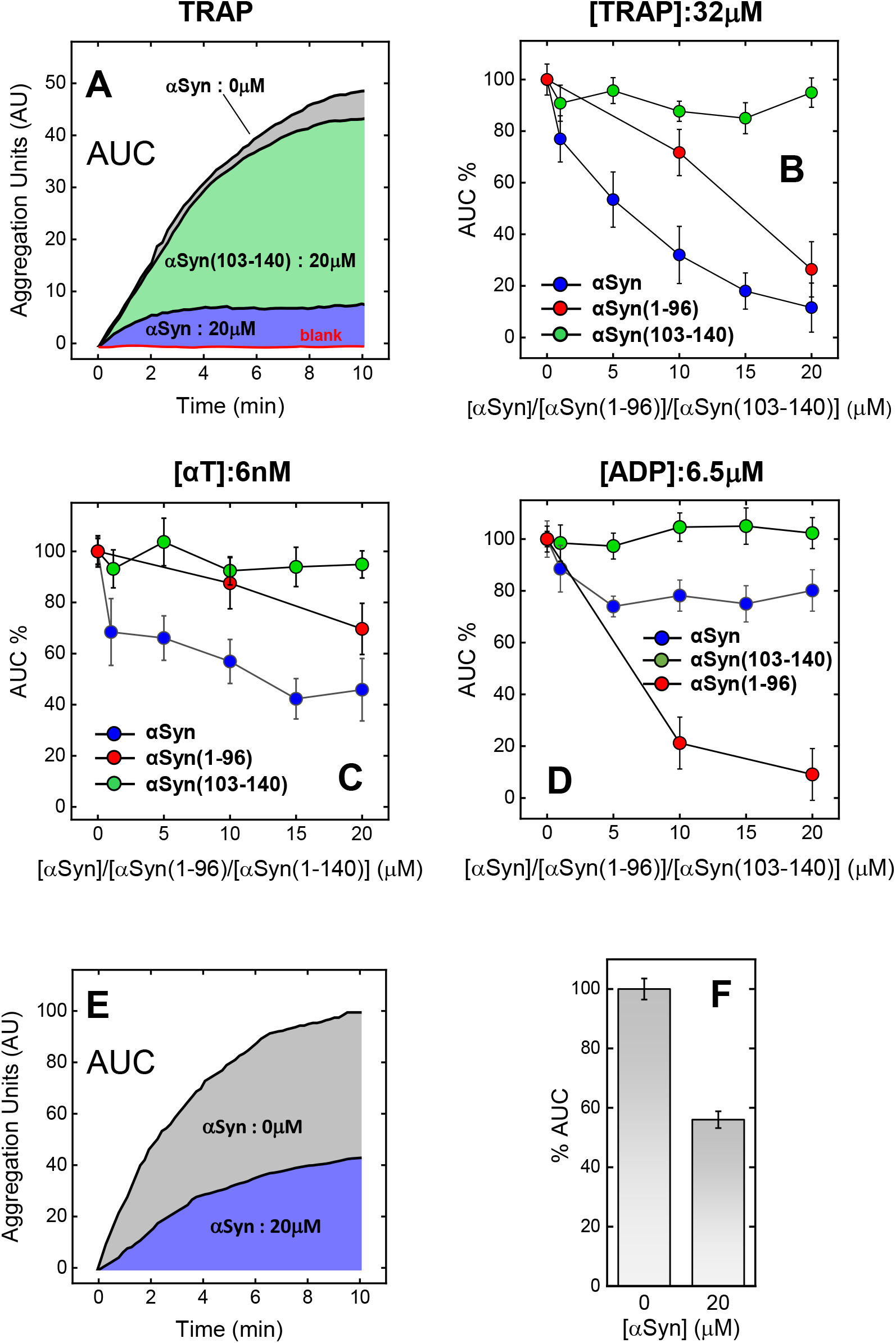
Effect of αSyn, αSyn(103-140) and αSyn(103-140) on the aggregation of platelets induced by TRAP, αT or ADP. (**A**) Representative impedance aggregometry curves of platelets aggregation induced by TRAP on whole blood in the absence (grey area) and presence of 20 μM αSyn (blue area) or αSyn(103-140) (green area). (**B-D**) Impedance aggregometry analysis of platelet agglutination induced by 32 μM TRAP6 (**B**), 6 nM αT (**C**), and 6.5 μM ADP (**D**) on whole blood at 37°C at increasing concentrations of full-length αSyn 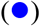 αSyn(103-140) 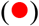 or αSyn(103-140) 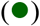. At each αSyn concentration, the Area Under the Curve (%AUC) was calculated relative to the value determined in the absence of αSyn (AUC_0_). Whole blood samples from healthy donors (160.000 platelets/μl) were incubated with solutions of αSyn species in HBS. Platelets agglutination was started by adding 20 μl of TRAP6 or ADP stock solutions. When the effect of αSyn on αT-induce aggregation was measured, protease solutions were pre-incubated with increasing concentrations of αSyn species and then added to blood samples. (**E, F**) αT-induced aggregation of isolated platelets in the presence of αSyn. (**E**) Representative impedance aggregometry curves obtained after addition of αT (6 nM) to washed human platelets (200·10^3^/μl) at 37°C in the absence (grey) and presence of 20 μM αSyn (blue). (**F**) Histogram plot of the platelets anti-aggregating effect of αSyn, as obtained for the data in panel **E**. For all measurements, each AUC value is the average of single determinations on blood samples from three healthy donors. The error bars correspond to the standard deviation. For further details, see Methods.

MEA is a fast and specific platelet-function-testing method (35), widely used in basic hematologic research and clinical testing, and enables to selectively measure aggregation of platelets, not only in isolation, as with classical light transmission aggregometry, but also in whole blood, which is the physiological environment where platelet function takes place *in vivo* (36). Indeed, the presence of erythrocytes and leukocytes in whole blood has been shown to influence platelet aggregation (36). The physical basis of MEA relays on the increase of impedance (i.e. the electric resistance to the passage of alternate current in a medium between two platinum electrodes) which is caused by sticking and subsequent intercellular adhesion of activated platelets onto the electrodes (35,36). Quiescent platelets stick to the electrodes and self-organize in cell monolayers. At this stage, platelets-electrode interaction does not increase the impedance signal. Only after addition of a platelet aggregant (e.g. αT, TRAP6, or ADP), activated platelets tightly adhere to the pre-existing monolayers on the electrodes, thus increasing blood electric impedance. For αT-induced platelet activation, fibrin fibers, generated after addition of αT to whole blood, might interact with activated platelets and further increase the impedance signal. To get a quantitative estimate of platelets aggregation, the time-dependent change of blood impedance is expressed as relative Aggregation Units (AU), where 8 AU approximately correspond to 1 Ohm (Ω). Integration of AU over time gives the value of the Area Under the Curve (AUC), which is taken as the best parameter of platelet function in MEA analysis (36).

Notably, αSyn has been reported to interact with metal surfaces (e.g. stainless steel and gold) (37). This might in principle hinder binding of platelets to the electrode surface, thus reducing stimulated platelets aggregation in MEA measurements, compared to controls, with a resulting apparent (artefactual) higher antiaggregating effect of αSyn. However, this possibility can be safely ruled out considering that: i) αSyn binds to metal surfaces in the fibrillar state, not in the monomeric state (37); ii) the lag phase of αSyn fibril formation, even in a crowded environment, is much longer (~3 h) than the time scale of MEA analysis (i.e. 6-10 min) (11); iii) finally, the tendency of αSyn oligomers/polymers to stick on metal surfaces is 2-3 times lower than that measured for albumin (37), which is even much more concentrated in the blood (260-380 μM) than the maximal αSyn concentration explored in this study (20 μM).

The data in **Fig. 2A-D** show that addition of full-length αSyn to whole blood samples variably reduced, in a concentration-dependent manner, the platelet aggregation potential (%AUC) of all activators tested, where the strongest inhibitory effect was observed at 20 μM αSyn with TRAP6 activation (~90%), followed by αT (~50%) and ADP (~20%). An IC_50_ value of 8.6±2.5 μM was estimated for the inhibition of αSyn on TRAP6-induced activation. As a negative control, addition of αSyn to whole blood, in the absence of TRAP6, does not induce any increase of AUC (**Fig. 2A**). To rule out the possibility that other cellular and soluble components, present in whole blood, might interfere with platelet aggregation, MEA analysis was also performed with platelets rich plasma (PRP) (**Fig. 2E,F**) Our data indicate that αSyn (20 μM) impairs αT-induced platelet agglutination by about **50%**, consistent with data obtained on whole blood (**Fig. 2C**). Noteworthy, the fact that αSyn impairs platelets aggregation, also induced by TRAP6 binding to PAR1, suggests that αSyn can directly interfere with PAR1 recognition, independently of αT-induced proteolytic activation.

As reported in the Introduction, αSyn is a small, disordered and acidic protein (pI 4.7) displaying a highly asymmetric charge distribution, with the N-terminal region strongly electropositive and the C-terminal tail strongly electronegative. Hence, to dissect the platelet anti-aggregating effect of αSyn, MEA measurements were also carried out with recombinant 6xHis-αSyn(1-96) (pI 9.4) and the synthetic αSyn(103-140) peptide (pI: 3.1) (**Fig. 2B-D**). Interestingly, the positively charged 6xHis-αSyn(1-96) retained the inhibitory effect of full-length αSyn towards platelets activation triggered by TRAP6, while the inhibition of activation by ADP was even stronger than that observed with αSyn. At variance, 6xHis-αSyn(1-96) impaired αT-induced platelet activation to a lower extent compared to αSyn. Noteworthy, and conversely to αSyn and 6xHis-αSyn(1-96), the negatively charged αSyn(103-140) displayed weak, if any, inhibitory potency in all platelet anti-aggregating assays tested (**Fig. 2B-D**).

### Effect of αSyn on fibrin generation

With the aim to investigate the possible effect of αSyn on fibrin generation, another key process in blood coagulation, the time-course change of turbidity (τ) of a fibrinogen (Fb) solution was monitored after addition of αT, in the presence of increasing αSyn concentrations (**Fig. 3**).

**Figure 3.**
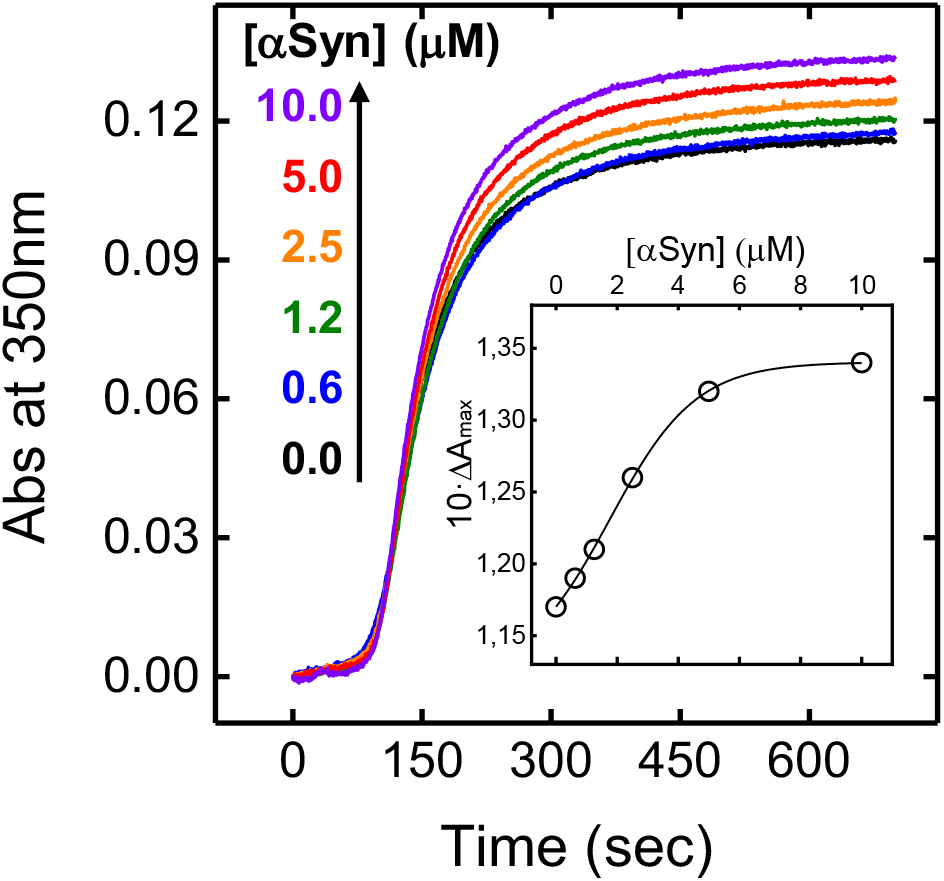
Effect of αSyn on αT-mediated fibrin generation. Turbidimetric analysis of fibrin generation induced by αT on purified fibrinogen at increasing αSyn concentrations, as indicated. To a desalted fibrinogen solution (440 nM, 800 μl) in HBS at 37°C, containing 0.1% PEG-8000, was added αT (1 nM, final concentration) pre-incubated with increasing [αSyn] and the time-course generation of fibrin was monitored by recording the absorbance increase of the solution at 350 nm. From each clotting curve, the values of t_c_, and ΔA_max_ were extracted. (**Inset**) Plot of ΔA_max_ vs. αSyn concentration.

Notably, τ is the decrease in the intensity of transmitted light at 350 nm, caused by the scattering of light due to fibrin formation, and is measured as the apparent absorbance increase of a Fb solution. Typically, a fibrin clotting curve (i.e. the time-course increase of τ) displays a sigmoidal shape, with i) a lag phase, corresponding to the time necessary for the longitudinal elongation of protofibrils; ii) a linear rise of the turbidity signal, resulting from lateral aggregation of protofibrils above a certain threshold length; and iii) a plateau, when most of protofibrils have been transformed into fibers, which then branch and assembly into the final fibrin network (38). From each clotting curve, the values of A_max_ and t_c_ can be extracted, where A_max_ is the maximum A_350nm_ value while t_c_ is the clotting time, defined as the time required for longitudinal elongation of fibrin fibers (see the legend to **Fig. 3**). Noteworthy, the value of A_max_ provides a key geometric parameter of fibrin structure, as it is proportional to the square of the average diameter of the fibers (38).

The data shown in **Fig. 3** indicate that, in the presence of αSyn, t_c_ remains essentially constant (109 ± 5 sec), whereas A_max_ progressively increases up to ~17% at 10 μM αSyn (**Fig. 3, Inset**). These results suggest that αSyn does not alter the lag phase of fibrin formation, when longitudinal fibrin polymerization occurs, while perturbing lateral aggregation to induce the formation of thicker fibers. These results can be explained considering that both longitudinal polymerization and lateral aggregation are variably influenced by charge-charge interactions, through which the “knobs” that are produced after fibrinopeptides release on one fibrin monomer interact with the “holes” present on another monomer (39). Therefore, it is not surprising that a highly charged protein like αSyn can interfere with fibrin structure. Noteworthy, a similar increase of A_max_ was observed in fibrin generation experiments at high ionic strength or in the presence of positively charged proteins, such as platelet factor-4 (pI: 8.9) (38).

### Effect of αSyn on αT-catalyzed substrate hydrolysis

αSyn was incubated with αT and then the protease hydrolytic activity was measured with specific substrates, including (D)-Phe-Pip-Arg-*p*-nitroanilide (S2238), fibrinogen (Fb), and the synthetic peptide PAR1(38–60) encompassing PAR1 activation domain.

The kinetics of *p*-nitroaniline release (**Fig. 4A**) clearly indicate that αSyn, up to the highest concentration explored (15 μM), does not appreciably affect the rate of S2238 hydrolysis and, furthermore, identical results were obtained with the same concentration of αSyn(103-140) (not shown). Likewise, the release of fibrinopeptides (i.e. FpA and FpB) from a fibrinogen solution was not affected by 15 μM αSyn, as documented by the invariance of the specificity constants (k_cat_/K_m_), extracted from the data in **Fig. 4B**. Noteworthy, αSyn (15 μM) was found to reduce by 2-fold the efficiency of PAR1(38–60) hydrolysis by αT (**Fig. 4C**), where the latter peptide reproduces the substrate binding properties of PAR1 extracellular domain on platelets, as it contains both the exosite-1 binding sequence for αT and the scissile bond Arg^41^-Ser^42^.

**Figure 4.**
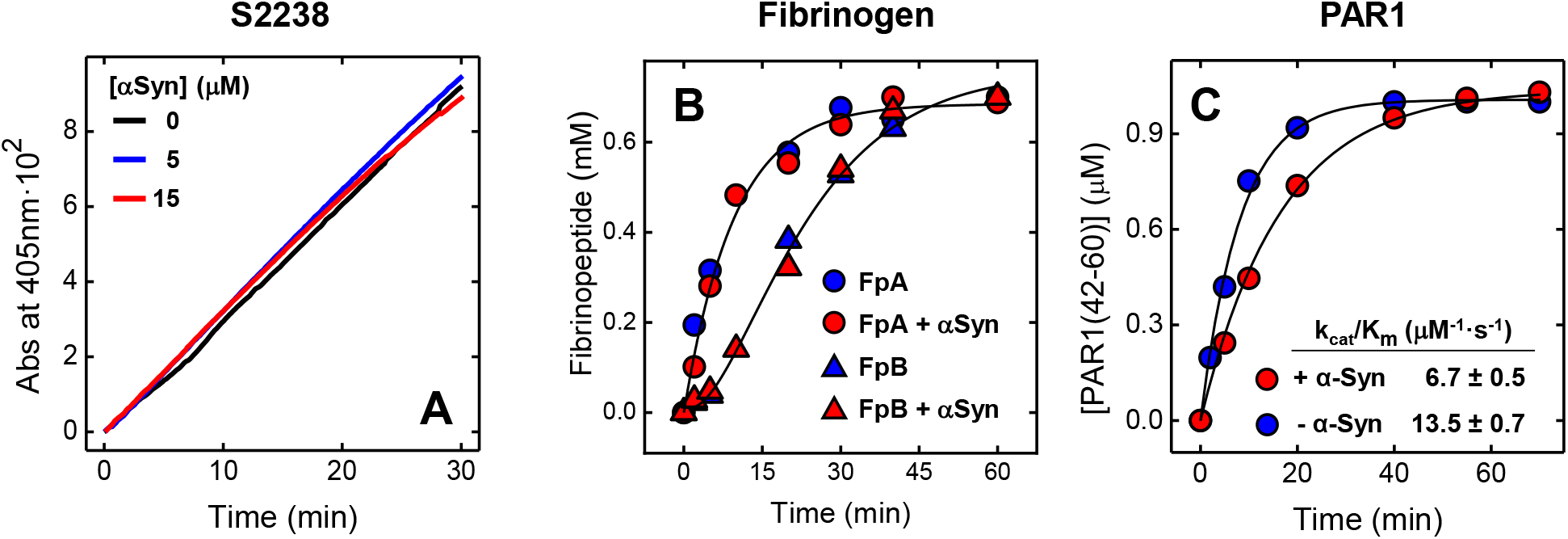
Effect of αSyn on αT-mediated hydrolysis of S2238, Fb and PAR1(38–60). (**A**) αT-catalyzed hydrolysis of the chromogenic substrate S2238 in the presence of increasing αSyn concentrations. The hydrolytic activity was determined at 37°C in HBS by measuring the release of *p*-nitroaniline (*p*NA) at 405 nm. (**B**) Release of FpA and FpB from desalted fibrinogen (0.35 μM) by αT (300 pM), in the absence or presence of αSyn (15 μM). Measurements were carried out at 37°C in HBS and quantified by RP-HPLC (see Methods). Interpolation of the data points with eq. 1 and 2, yielded the apparent specificity constants (k_cat_/K_m_) of FpA and FpB release in the absence (k_cat,A_/K_m,A_ = 7.2±0.9 μM^−1^·s^−1^; k_cat,B_/K_m,B_ = 5.7±0.6 μM^−1^·s^−1^) and presence (k_cat,A_/K_m,A_ = 4.7±2.2 μM^−1^·s^−1^; k_cat,B_/K_m,B_ = 4.6±2.1 μM^−1^·s^−1^) of 15 μM αSyn. (**C**) Cleavage of PAR1(38–60) by αT in the absence 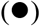 and presence 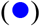 of αSyn (15 μM). The cleavage of PAR1(38–60) peptide (1 μM) by αT (150 pM) was carried out at 25°C in TBS and the time course of PAR1(42–60) fragment release quantified by RP-HPLC. The data points were fitted with eq. 3, to yield the values of k_cat_/K_m_ in the absence and presence of αSyn.

### Probing αSyn-αT interaction by fluorescence spectroscopy and surface plasmon resonance (SPR)

The interaction of αSyn with αT was monitored by steady state fluorescence change and SPR, two orthogonal techniques exploiting different physico-chemical observables. αSyn-αT interaction was also probed by isothermal titration calorimetry, which however yielded a very low heat exchange, too small to obtain a reliable equilibrium affinity constant (not shown).

A first evidence of αSyn-αT complex formation came from fluorescence emission spectra **(Fig. 5A)**, obtained after excitation at 295 nm, indicating that addition of αSyn (20 μM, final concentration) to a αT solution (70 nM, final concentration) reduced by **~10%** the fluorescence intensity of the solution, compared to the theoretical sum-spectrum of both isolated αT and αSyn at the same concentrations, without appreciably altering the wavelength of maximum emission (λ_max_), i.e. 334 nm. The fluorescence change associated with αSyn-αT coupling is likely caused by changes in the environment of some Trp-residues in αT structure. Indeed, αT contains nine tryptophan fluorophores, whereas αSyn has only four Tyr-residues, which do not (or only negligibly) absorb at 295 nm and, therefore, are not expected to contribute to the emitted fluorescence. A quantitative estimate of αSyn-αT interaction was obtained by recording the decrease of αT fluorescence at λ_max_ at increasing αSyn concentrations (**Fig. 5B**). Interpolation of the data points with eq. 4, describing a stoichiometric 1:1 binding model, yielded an equilibrium dissociation constant (K_d_) of 0.96 μM.

**Figure 5.**
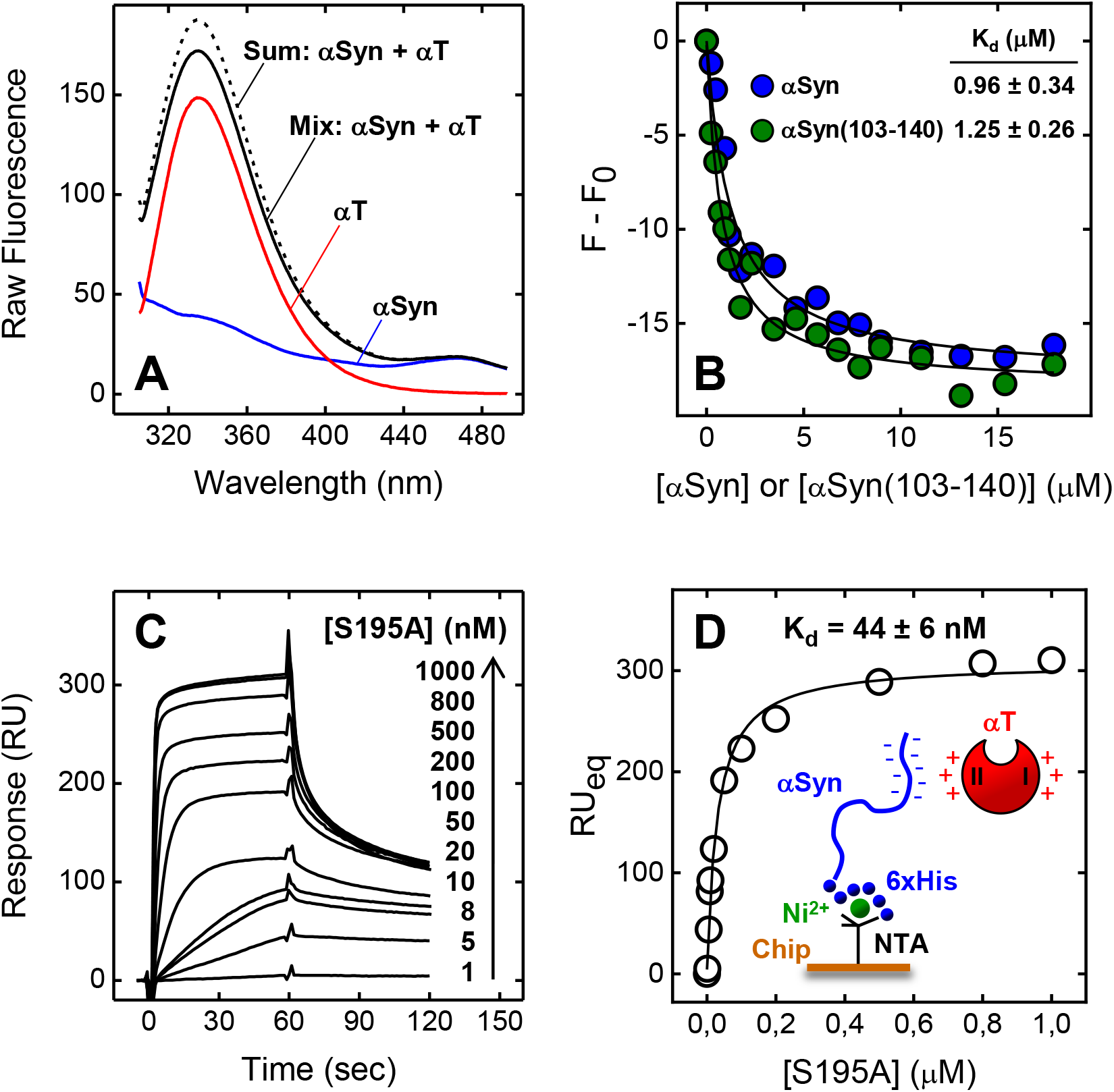
Probing αT-αSyn interaction by fluorescence spectroscopy and surface plasmon resonance. (**A**) Fluorescence spectra of isolated αSyn (20 μM, **–**) and αT (70 nM, **–**), and of the experimental complex containing the interacting proteins at the corresponding concentrations (**–**). For comparison, the theoretical sum spectrum (---) is also reported. Emission spectra were recorded in HBS at 37°C, after excitation at 295 nm, and subtracted for the corresponding baseline. (**B**) Fluorescence binding measurements of αSyn interaction with αT. To a solution of αT (70 nM) in HBS at 37°C were added aliquots (2-20 μl) of full-length αSyn 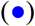 and αSyn(103-140) 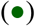. The samples were excited at 295 nm, and the emission intensity was recorded at 334 nm. Each spectrum was subtracted for the contribution of αSyn alone at the corresponding concentration and expressed as F - F_0_, where F and F_0_ are the fluorescence intensity in the presence or absence of αSyn derivatives. The data points were interpolated with eq. 4, yielding the K_d_ value as indicated. (**C, D**) SPR analysis of αT binding to immobilized αSyn. Recombinant wild-type 6xHis-αSyn was immobilized onto a Ni^2+^-NTA sensor chip and increasing concentrations of S195A thrombin mutant were injected in the mobile phase. (**C**) Sensograms relative to S195A binding. (**D**) Plot of RU_max_ vs. S195A concentration (∘). Fitting of data points with eq. 6 yielded the K_d_ value for the synuclein-thrombin complex, as indicated. All SPR measurements were carried out at 37°C in HBS-EP^+^.

Thereafter, SPR measurements were carried out by immobilizing the N-terminally 6xHis-tagged αSyn onto a Ni^2+^/nitrilotriacetate sensor chip, *via* non-covalent chelation, and injecting incremental concentrations of S195A in the mobile phase (**Fig. 5C,D**). The catalytically inactive S195A thrombin mutant was used, as active αT was shown to cleave the fused 6xHis-αSyn at Lys6, but not the untagged wild-type αSyn 1-140 (**Supplementary Fig. S2**). SPR data were analyzed according to the one-site binding model (eq. 5), yielding a K_d_ of 44 nM. Notably, the affinity of αT for immobilized αSyn was >20-fold higher than that estimated by fluorescence binding experiments. Overestimation of binding strength in protein-protein interactions is frequent in SPR measurements, compared to other spectroscopic techniques (e.g. fluorescence), and it is inherently associated to the beneficial lower loss of binding entropy (ΔS_b_) occurring in a biphasic interacting system like SPR, where one of the two partners is immobilized on the sensor chip (40,41). This is especially true for an intrinsically unfolded protein like αSyn, which becomes “more ordered” after immobilization on the sensor chip, with a resulting favorable reduction of ΔS_b_, compared to the solution phase where both interacting partners are “free” in solution and therefore undergo a larger (unfavorable) entropy loss of binding (40).

Altogether, both fluorescence and SPR measurements concurrently indicate that αSyn binds to αT with a moderate to high affinity, depending on the binding system investigated.

### Molecular mapping of αSyn-αT interaction

To identify the region of αSyn responsible of αT binding, we measured the affinity of αSyn(103-140) for αT by steady state fluorescence spectroscopy (**Fig. 5B**). Our data indicate that αSyn(103-140) binds to αT with an affinity very similar (K_d_ = 1.25 μM) to that of full-length αSyn (K_d_ = 0.96 μM), suggesting that the negatively charged C-terminal tail of αSyn is the protein binding epitope for αT.

Next, we mapped the sites on αT structure that are involved in the interaction with αSyn. The active site and two positively charged patches, namely exosites 1 and 2, are the hot spots on αT responsible for the recognition of most physiological substrates and inhibitors (42–44). The role played by these regions in binding to αSyn was assessed using “the site-specific perturbation approach”, earlier exploited in our laboratory in the study of αT interactions (41,45-48). Here, the effect of saturating concentrations of αSyn on the affinity of selected ligands, which are known to specifically bind to a given target site on αT structure (e.g. the active site, exosite 1 or 2), was measured. The decrease in binding strength for a given site-specific ligand was taken as a strong indication that the target site is involved in binding to αSyn. Furthermore, we used thrombin derivatives in which one of the key binding sites was selectively perturbed by proteolytic nicking [i.e. β_T_-thrombin (β_T_T)] or intramolecular masking [i.e. prothrombin (ProT)]. Once again, a drop in binding affinity provides evidence for the involvement of the perturbed region in the interaction with αSyn.

### Active Site

To check whether the active site region of αT is involved in αSyn binding, the affinity of ligands/inhibitors of incremental size and mapping different αT subsites [i.e. *p*-aminobenzamidine (PABA), the chromogenic substrate S2238, and hirudin fragment 1-47] was measured in the absence and presence of 20 μM αSyn. PABA, a small fluorescent inhibitor of chymotrysin-like proteases, interacts with the substrate primary specificity site (S1) of αT and, after binding, emits more intensely at 375 nm. The αT-specific substrate S2238 extensively interacts with the protease recognition sites (49) and its affinity for the catalytically inactive thrombin mutant S195A was determined by fluorescence resonance energy transfer measurements, where the *p*-nitroanilide moiety of S2238 functions as the energy acceptor while the Trp-residues of S195A act as the energy donors (45). Hirudin fragment 1-47, i.e. Hir(1–47), is a potent thrombin inhibitor covering the active site and the loop regions nearby and, upon binding to αT, induces an increase of the protease fluorescence emission (50).

The data in **Fig. 6** indicate that αSyn only marginally alters the affinity of all active-site specific ligands tested, suggesting that the αT catalytic site is not significantly involved in αSyn binding. These data are consistent with the observation that αSyn has no effect on the hydrolysis rate of S2238 by αT (**Fig. 4**).

**Figure 6.**
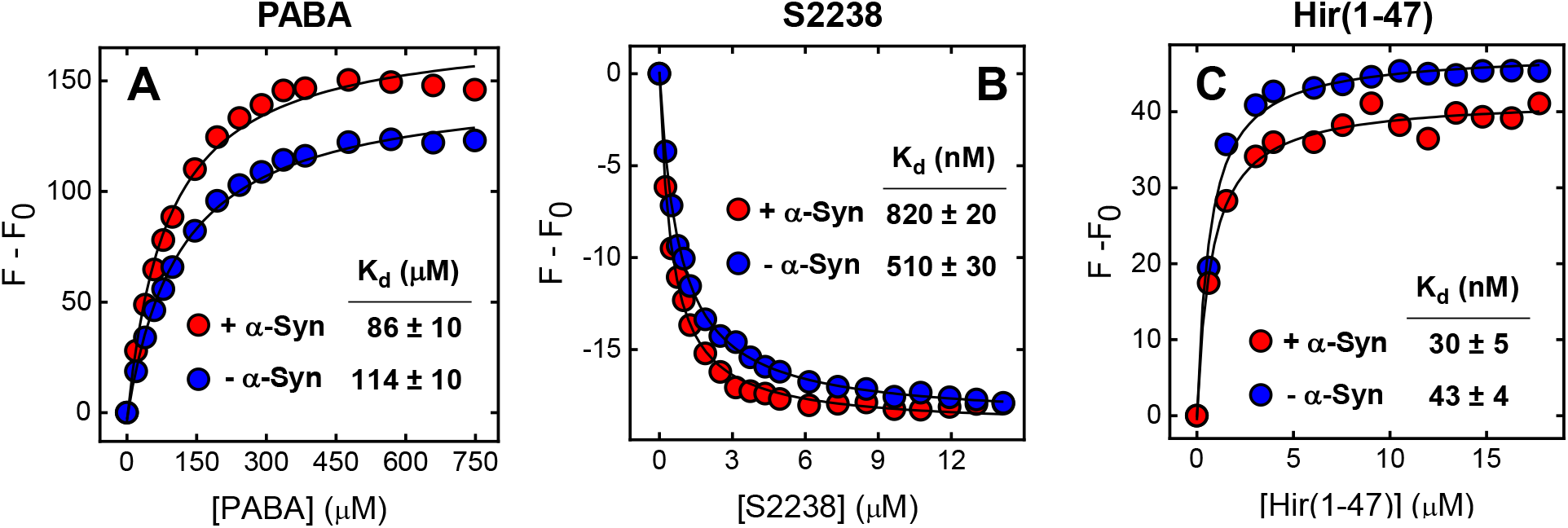
Probing the role of thrombin active site in αSyn-αT interaction. Effect of αSyn on the affinity of the active-site ligands PABA (**A**), S2238 (**B**), or Hir(1–47) (**C**) for thrombin. Fluorescence binding measurements were carried out in HBS at 37°C by adding increasing ligand concentrations to thrombin solutions, in the absence or presence of 20 μM αSyn. For the binding of PABA, samples (40 nM) were excited at 336nm, the fluorescence intensity of the ligand was recorded at 375 nm and corrected for inner filter effect. With S2238 and Hir(1–47), protein samples (50 nM and 70 nM, respectively) were excited at 295 nm, while thrombin fluorescence was recorded at 334 nm. When the binding of S2238 to thrombin was being studied, the inactive S195A mutant was used. The data points relative to the binding of PABA and S2338 were interpolated with eq. 4, describing a single-site interaction model, while the data for the binding of Hir(1–47) were fitted with eq. 5, describing a tight-binding model. After interpolation, the K_d_ values were obtained as fitting parameters, as indicated.

*Exosites 1 and 2*. The involvement of αT exosites in binding to αSyn was probed by fluorescence binding measurements, measuring the affinity of exosite-specific ligands for the protease in the absence and presence of αSyn or αSyn(103-140). Hirugen (i.e. the C-terminal peptide 54-65 of hirudin HV1) was selected as a safe exosite-1 binder (41,43), while the fibrinogen γ’-peptide (i.e. the C-terminal peptide 408-427 of fibrinogen elongated γ-chain splice variant) was used as a specific exosite-2 ligand (51,52). Full-length αSyn and αSyn(103-140) decreased the affinity of the γ’-peptide for αT by about 3- and 4-fold (**Fig. 7C**,**D**), respectively, but they were not able to reduce the affinity of hirugen for exosite 1 (**Fig. 7A**,**B**). A similar conclusion was drawn from displacement experiments (41,46), showing that neither αSyn nor αSyn(103-140) were able to displace N^α^-fluoresceinated hirugen, [F]-hirugen, from αT exosite 1 (**Fig. 7E**).

**Figure 7.**
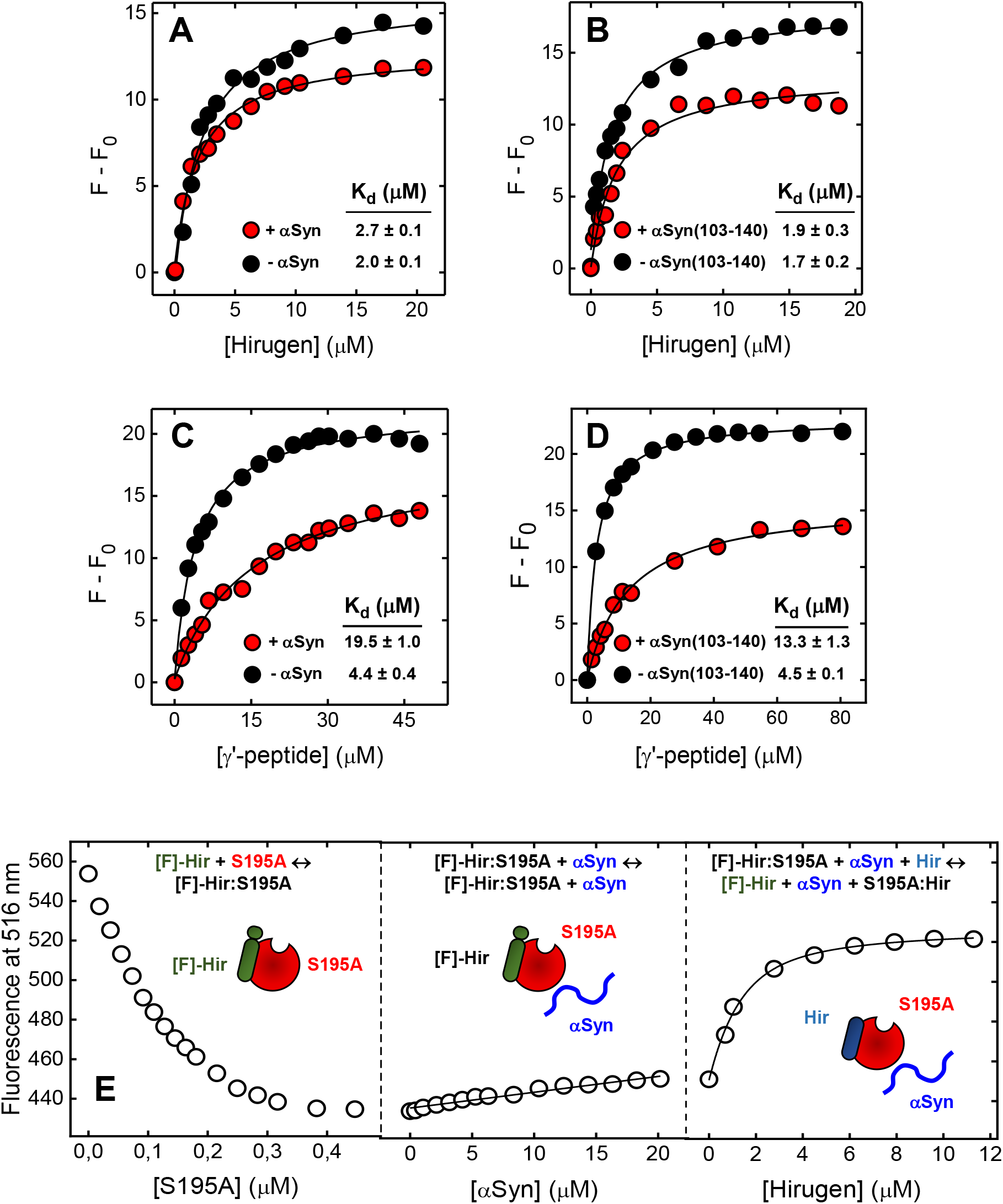
Probing the role of thrombin exosites in αSyn-αT interaction. (**A, B**) The role of exosite 1. Effect of αSyn and αSyn(103-140) on the affinity of hirugen for thrombin exosite-1. Fluorescence measurements of hirugen binding to αT (70 nM) in the absence or presence of saturating concentrations (20 μM) of full-length αSyn (**A**) or αSyn(103-140) (**B**). (**C, D**) The role of exosite 2. Effect of αSyn and αSyn(103-140) on the affinity of γ′-peptide for thrombin exosite-2. Fluorescence measurements of γ′-peptide binding to αT (70 nM) in the absence or presence of saturating concentrations (20 μM) of full-length αSyn (**C**) or αSyn(103-140) (**D**). Protein samples in HBS at 37°C were excited at 295 nm and the fluorescence intensity was recorded at 334 nm, after baseline subtraction. The data points were interpolated with eq. 4 to obtain the K_d_ values, as indicated. (**E**) Displacement of [F]-hirugen from αT exosite −1 by αSyn. *Left panel*, Binding of S195A to [F]-hirugen (60 nM). The data points were interpolated with eq. 5, yielding a K_d_ of 30 ± 8nM. *Middle panel*, Displacement of [F]-hirugen from the complex with S195A by incremental concentrations of αSyn. A moderate increase of fluorescence (~17%) was observed, suggestive of only a partial displacement of [F]-hirugen by αSyn. *Right panel*, Displacement of [F]-hirugen from S195A by incremental concentrations of unlabelled hirugen. A marked increase of fluorescence (~65%) was measured as the result of complete displacement of residual [F]-hirugen from αT exosite-1. Samples in HBS, containing 0.1% PEG-8000, were excited at 25°C at 492 nm and the fluorescence of [F]-hirugen, bound to or released from S195A, was recorded at 516 nm.

The role of αT exosites in αT-αSyn interaction was further investigated by SPR, whereby the binding strength of thrombin species (i.e. β_T_T and ProT), having the exosites variably compromised, to immobilized 6xHis-αSyn was measured (**Fig. 8**). β_T_T results from proteolytic nicking of mature αT by trypsin at the peptide bond Arg77a-Asn78, leading to disruption of exosite-1, whereas the active site and exosite 2 retain the structural and functional properties of the corresponding regions in mature αT (49) (**Fig. 8A**). ProT is the physiological zymogen precursor of αT and, compared to αT, major structural perturbations occur in the Na^+^-binding site, the activation domain and the insertion loops surrounding the catalytic cleft (47). Importantly, exosite-1 structure seems to be only slightly altered in the zymogen structure, whereas the reactivity of exosite 2 is completely abolished because of intramolecular tight binding of the zymogen kringle-2 domain (**Fig. 8A**). Analysis of binding data (**Fig. 8B**) shows that disruption of exosite 1, as in β_T_T, reduces the affinity for 6xHis-αSyn by only 1.5-fold, whereas masking of exosite 2, as in ProT, leads to a dramatic drop in binding strength by 37-fold. These findings highlight the role of exosite 2 in αSyn-αT interaction, as masking/perturbation of exosite 2 results in a drop of the affinity of αSyn for thrombin, whereas masking/perturbation of exosite 1 is essentially not relevant.

**Figure 8.**
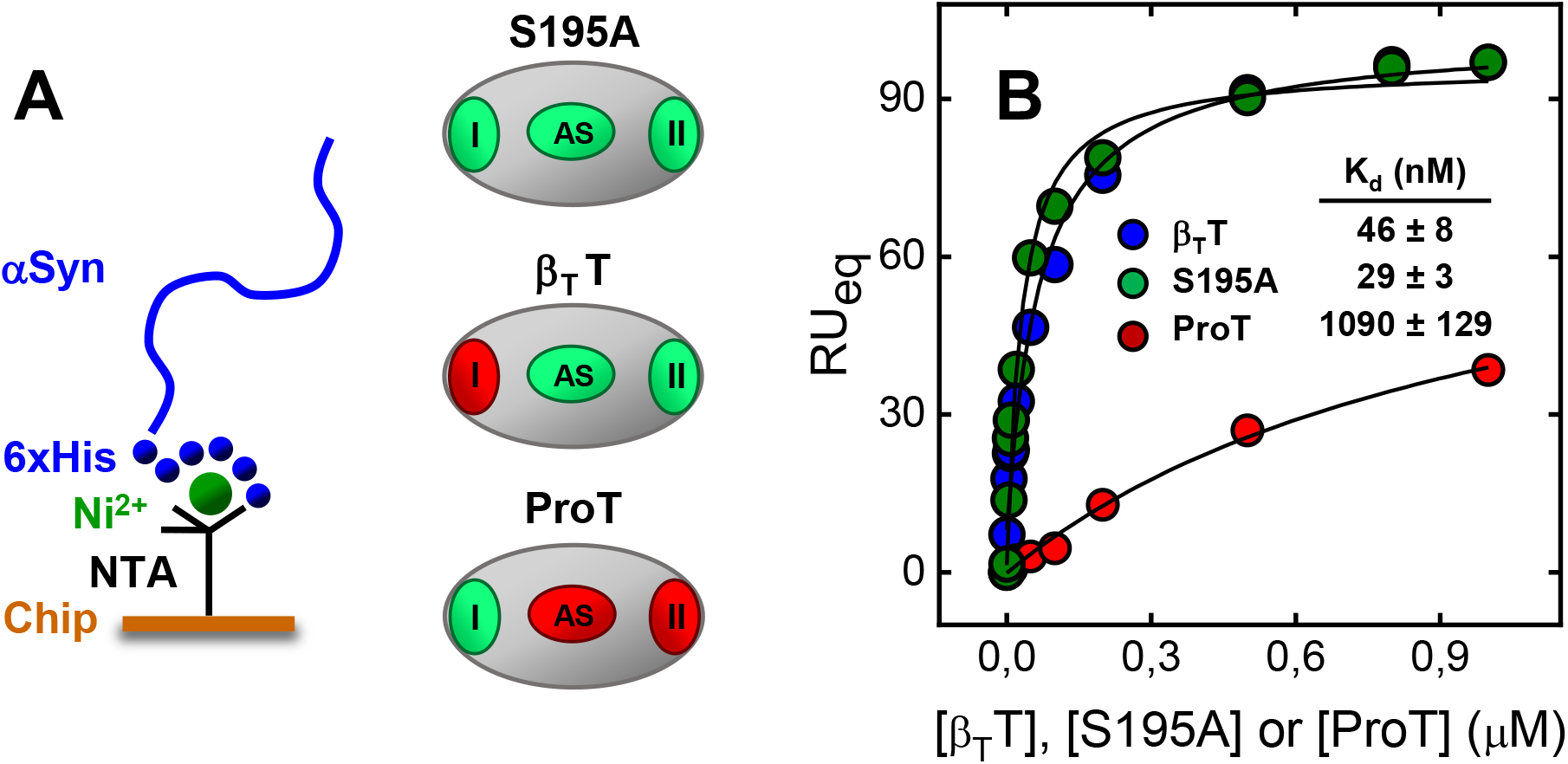
Effect of selective perturbation of thrombin exosites on the affinity for αSyn. (**A**) Schematic representation of the protease-domain of rS195A thrombin mutant, β_T_-thrombin (β_T_T), and prothrombin (ProT). The active site (AS) and exosite-1 (I) and exosite-2 (II) are colored according to the conformational/functional state they assume in the different thrombin derivatives, compared to αT (see text); *green*: unperturbed or only slightly perturbed; *red*: heavily perturbed. (**B**) SPR analysis of the binding of S195A, β_T_T, and ProT to 6xHis-αSyn immobilized on a Ni^2+^-NTA sensor chip. The values of RU_max_ were plotted *versus* the concentration of thrombin derivatives and the data points were interpolated with eq. 6, yielding the corresponding K_d_ values as indicated. Measurements were carried out at 37°C in HBS-EP^+^, pH 7.4.

Altogether, the results of molecular mapping experiments, conducted by fluorescence and SPR measurements on the αSyn-αT interacting system, concurrently indicate that αSyn uses its negative C-terminal tail 103-140 to interact with the electropositive exosite 2 of αT.

### Probing aSyn membrane localization in resting and activated platelets by fluorescence microscopy

Platelets activation is characterized by the exposure of anionic phosphatidylserine from the inner to the outer leaflet of the plasma membrane, providing a negative binding surface for coagulation factors complex assembly and activation during the burst phase of αT generation (28). αSyn is abundantly present in the cytoplasm of resting platelets, associated with the membrane of secretory α-granules and the cytoplasmatic leaflet of the plasma membrane (20,23,24). Furthermore, *in vitro* studies indicate that the positive NT drives binding of αSyn to negatively charged membranes (5), where it folds in an α-turn-α motif, whereas the negative CT fluctuates outside the plasma membrane and remains largely unfolded, due to intramolecular charge repulsion and the presence of five secondary structure-destabilizing prolines (**Fig. 1**). Nevertheless, direct experimental data on the extracellular localization of αSyn on activated platelets is still lacking.

To fill this gap, exogenous αSyn-GFP, i.e. recombinant αSyn fused at the C-terminal end with the green-fluorescent protein (GFP) (31,32), was added to resting and TRAP6 activated platelets and plasma membrane binding was studied by fluorescence microscopy. Furthermore, localization of endogenous αSyn, before and after TRAP6 stimulation, was studied by immunofluorescence microscopy using a mouse anti-human αSyn monoclonal primary antibody, namely α-synuclein(211) recognizing αSyn 121-125 sequence, and an Alexa Fluor 594-conjugated goat anti-mouse IgG, as a secondary antibody.

Time-lapse fluorescence microscopy images of seeded platelets (20×10^6^ platelets/well) indicate that αSyn-GFP preferentially adheres, in a concentration dependent manner (0-2.0 μM), to activated platelets, compared to resting platelets (**Fig. 9A**). This result was confirmed in a separate experiment, where platelets (16×10^6^ platelets/well) were fixed with paraformaldehyde, before and after TRAP6 stimulation. The micrographs in **Fig. 9B** indicate that αSyn preferentially localizes on the plasma membrane of TRAP6-activated platelets. Finally, immunofluorescence microscopy was used to detect the exposure of endogenous αSyn on the plasma membrane of resting and activated platelets (16×10^6^ platelets/well). Our data indicate that a basal expression of αSyn on the plasma membrane is present even in resting platelets and that this expression is substantially increased after stimulation with TRAP6 (**Fig. 9C**).

**Figure 9.**
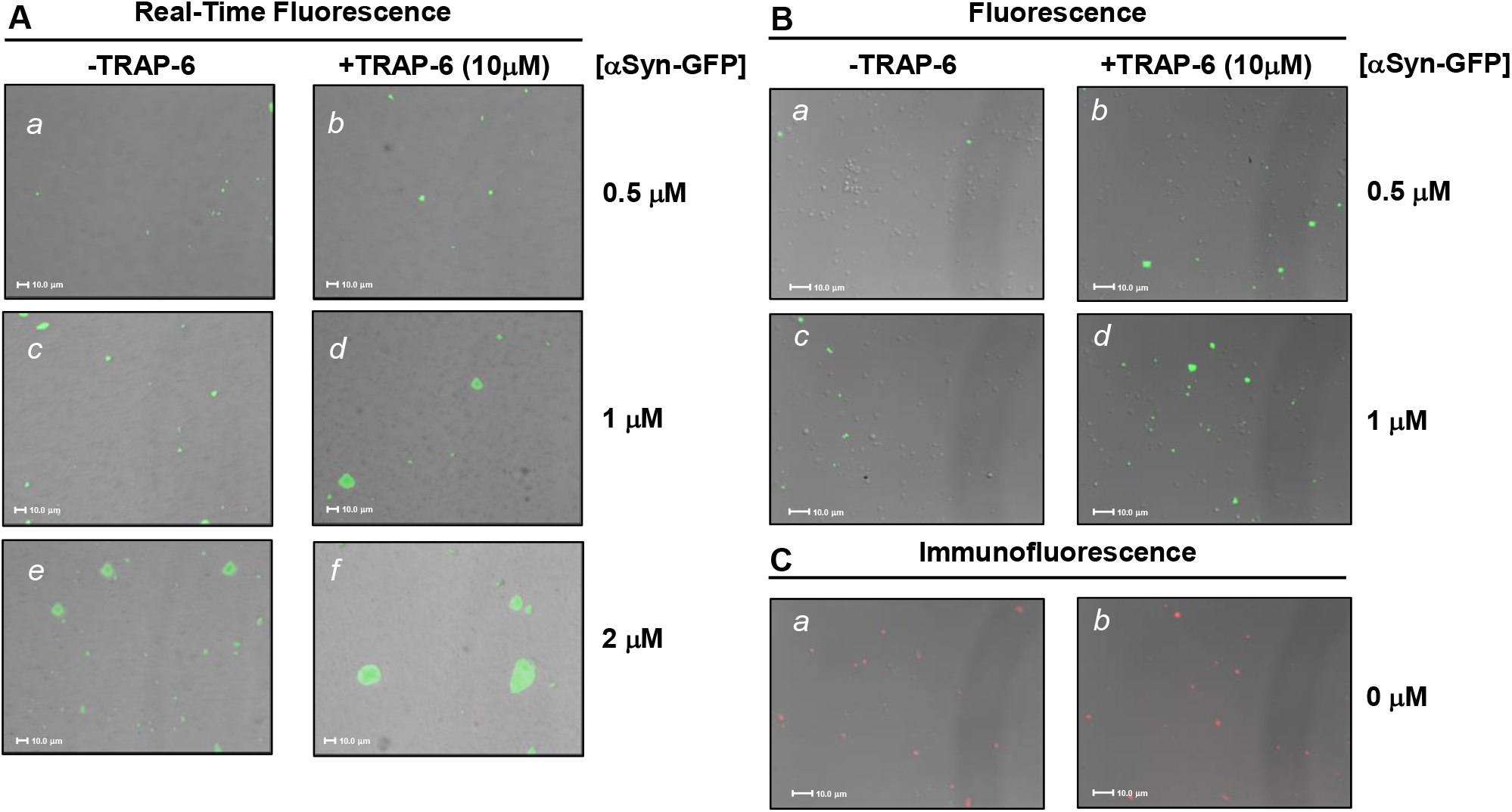
Fluorescence microscopy analysis of αSyn membrane localization in resting and activated platelets. (**A**) Time-lapse fluorescence microscopy. Isolated platelets were seeded (20×10^6^ platelets/well) in 24-well culture plates. After 24 h, increasing concentrations of αSyn-GFP (0, 0.5, 1.0, and 2.0 μM) in HBS were added to resting and TRAP6 (10µM) activated platelets, as indicated in panels *a*-*f*. After 30-min incubation at 37°C, under 5% CO_2_ flow, images were taken in real time. (**B**) Fluorescence microscopy. Seeded platelets, either resting or activated, (16×10^6^ platelets/well) were incubated for 24 h with αSyn-GFP (0.7 and 1.4 μM), washed twice with PBS and fixed for 20 min in 2% paraformaldehyde. (**C**) Immunofluorescence microscopy. Restring and activated platelets (not treated with αSyn-GFP) were seeded (16×10^6^ platelets/well), fixed with paraformaldehyde and incubated for 1 h at 37°C with mouse anti-human αSyn monoclonal antibody, namely α-synuclein(211), (5 μg/ml) followed by addition of a diluted (1:200) Alexa Fluor 594-conjugated goat anti-mouse IgG (red fluorescence). Both primary and secondary antibodies were diluted in PBS, containing 0.5% bovine serum albumin. Unspecific binding was assessed by incubating platelets with the secondary antibody alone, in the absence of the primary antibody. Final pictures resulted from merging differential interference contrast (DIC) and fluorescence images.

### Electrostatic properties of αSyn, αT, and platelet receptors

With the aim to rationalize on simple physical grounds the αT binding properties of αSyn and its platelets antiaggregating effect, we decided to investigate the electrostatic properties of αSyn and its interacting partners (i.e. αT and platelet receptors PAR1, PAR4 and P2Y_12_R), using the APBS software (53).

αSyn is a small acidic protein (pI 4.7) that, at physiological plasma pH, contains 16 positive and 25 negative charges (comprising the N- and C-termini), which account for about 30% of the protein amino acid content and actually impair αSyn to acquire a stable folded structure in solution (3). The unique electrostatic properties of the protein are self-evident from its amino acids sequence (**Fig. 1**), as the N-terminal (NT) region (1–60) contains an excess of seven positive charges, whereas the C-terminal (CT) region (96-140) is strongly negative with an excess of 12 negatively charged amino acids. The central NAC region 61-95, driving protein aggregation and fibrillogenesis, is highly hydrophobic and only slightly positive, with one positive charge in excess.

Even αT displays a non-uniform electrostatic potential, generated by a highly asymmetric distribution of positive and negative amino acids on the protease structure (**Fig. 10A**), whereby exosite 1 and 2 are strongly positive whereas the Na^+^-binding site and the active site region are negatively charged, with the catalytic pocket surrounded by a “negative ring” of Asp-and Glu-residues (47). Although both exosites display positive electrostatic potential, exosite 2 is more electropositive and ligand binding is mainly driven by “less specific” electrostatic complementarity, whereas exosite 1, beyond electrostatics, has “more specific” hydrophobic and stereochemical requirements for interaction (42–44).

**Figure 10.**
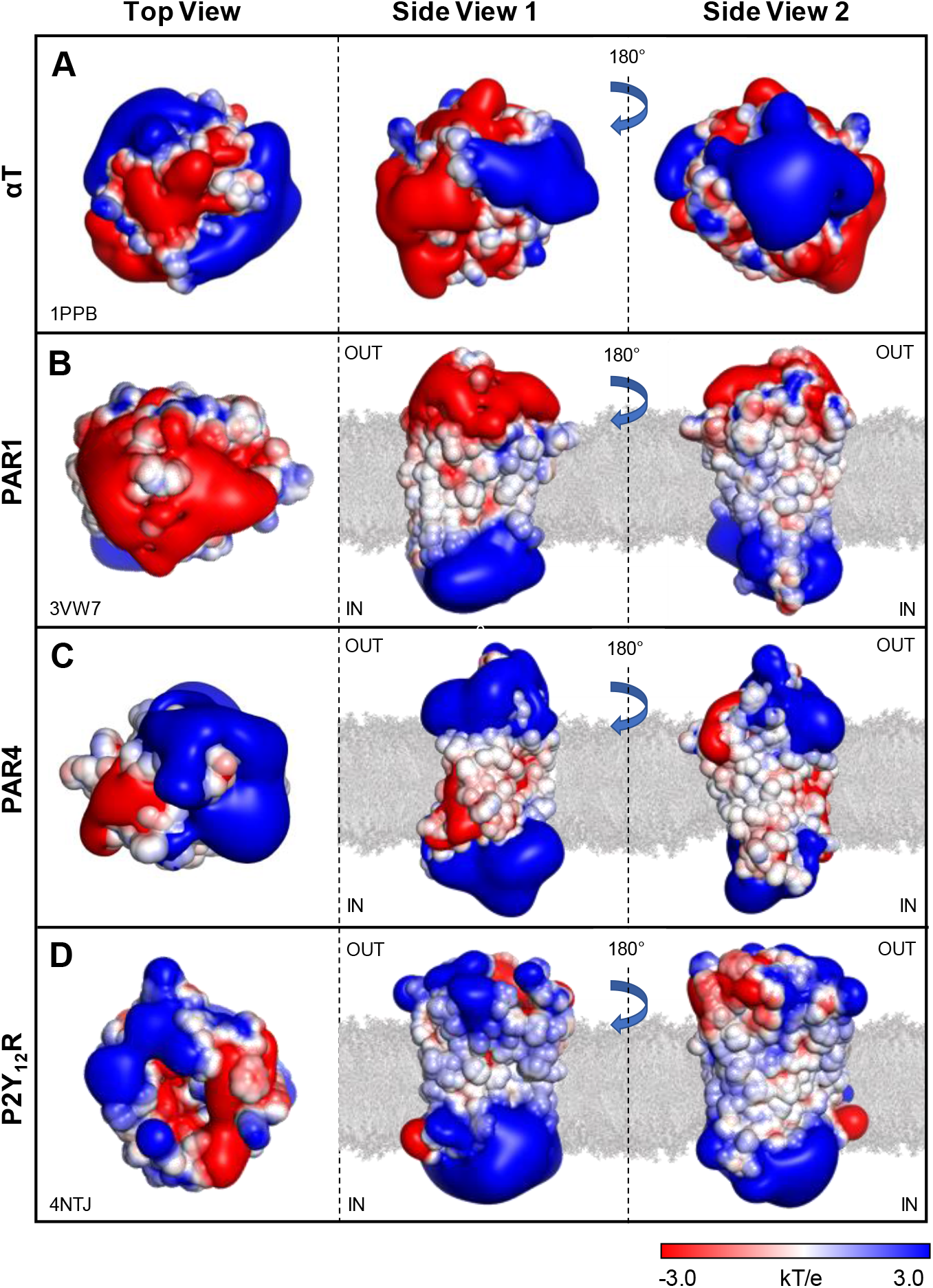
Surface electrostatic potential of αT (A), αT receptor PAR1 (B), and ADP receptor P2Y_12_R **(C)**. Calculations were carried out using the APBS software on the deposited structures of αT (1ppb), PAR1 (3vw7), P2Y_12_R (4ntj). PAR4 was modeled by homology on the PAR1 structure. Images were generated with PyMOL vs. 1.3 (DeLano Scientific, San Diego, CA, USA). The surface is coloured according to the electrostatic potential (blue, positive; red, negative) and expressed as kJ/(mol·q). Phospholipid double layer (grey) has been manually inserted. (**A**) αT displays an asymmetrical electrostatic potential, with a strongly negative active site (AS) region flanked by the two electropositive exosites (I and II). (**B**) The structure of PAR1, lacking the flexible exodomain A^22^-E^90^, displays a highly polarized charged distribution, with a strongly negative extracellular surface (OUT) and a positive intracellular region (IN). As expected, the transmembrane region in contact with the phospholipid apolar chains is essentially neutral. PAR1 contains a Na^+^-ion bound in the middle of the 7TM-helix bundle. The contribution of this ion was not taken into account during electrostatic calculations. (**C, D**) PAR4 and P2Y_12_R display a charge distribution similar to that of PAR1. However, contrary to PAR1, the extracellular region of P2Y_12_R and PAR4 is mainly electropositive with only some interspersed negative pots.

Electrostatic potential calculations (**Fig. 10**), carried out on the crystallographic structure of PAR1 (54) and P2Y_12_R (55) and on the modelled structure of PAR4 (see Methods), indicate that the cytoplasmatic region of all these receptors is always electropositive, whereas the extracellular region is strongly electronegative in PAR1 (**Fig. 10B**) and substantially electropositive in both PAR4 (**Fig. 10C**) and P2Y_12_R (**Fig. 10D**), with only some interspersed negative spots. As expected, the transmembrane regions, in contact with the membrane phospholipids apolar chains, are essentially neutral in all the receptors investigated. Besides the different charge distribution, PAR1 differs from PAR4 and P2Y_12_R in the binding mode to agonist/antagonist molecules. Indeed, whereas PAR1 ligands mainly exploit receptor electrostatic complementarity and interact with superficial structures rather than penetrating deeply into the receptor core (54), ligand binding to P2Y_12_R (55) and PAR4 (56) is more “stereochemically demanding” and it is mainly driven by orientation-dependent interactions, e.g. van der Waals interactions and hydrogen bonds.

## Discussion

This study follows our previous work aimed at identifying novel biochemical pathways at the interface of thrombosis (41,48,57,58) and amyloidosis (59).

Using MEA analysis on whole blood samples and isolated platelets, here we have shown that αSyn primarily impairs activation of platelets induced by TRAP6 (up to ~90%) and αT (up to ~50%) and, to a minor extent, by ADP (up to ~20%) (**Fig. 2**). Using several different molecular probes (i.e. PABA, S2238, Hir(1–47), hirugen and [F]-hirugen, γ’-peptide, thrombin S195A mutant, β_T_T and ProT) and techniques (i.e. fluorescence and SPR) we have also demonstrated that αSyn preferentially binds to αT exosite 2 through its negative C-terminal tail αSyn(103-140), without appreciable involvement of exosite 1 and active site (**Figs. 5–8**). Neither fibrinopeptides release nor S2238 hydrolysis was influenced by αSyn (15 μM), while the efficiency of PAR1(38–60) cleavage was only moderately reduced by 2-fold (**Fig. 4**). Finally, fluorescence microscopy data indicate that resting platelets basally express endogenous αSyn on the plasma membrane and that this expression is enhanced after stimulation with TRAP6. Likewise, exogenous αSyn adheres in a concentration dependent manner and to a greater extent to activated compared to resting platelets (**Fig. 9**).

From the comparison of platelets antiaggregating effect of full-length αSyn with that of its constituent peptides αSyn(1-96) and αSyn(103-140) (**Fig. 2B-D**), it is possible to conclude that: i) the positively charged N-terminal region αSyn(1-96) is required for αSyn anti-aggregating activity; ii) the negative C-terminal region αSyn(103-140) *per se* does not have any direct anti-aggregating effect; iii) the presence of αSyn(103-140) has a variable effect, either positive or negative, on αSyn anti-aggregating function, depending on the specific functional assay tested.

### Electrostatic model of aSyn platelets antiaggregating activity

Electrostatic charge-charge interactions play a pivotal role in the biochemistry of blood coagulation, as exemplified by the reversal of membrane charge polarization following platelet activation (see above) (28) or by the proteolytic activation of zymogens in the coagulation cascade (49), whereby active site formation in the mature protease is driven by the energy released after formation of a high-energy salt bridge between the newly generated N-terminus and a conserved Asp-residue in the activation domain. More specifically, proteolytic activation of platelet PAR1 by αT entails coupling of the enzyme electropositive exosite 2 with the highly negative C-terminal tail (amino acids 268-282) of platelet GpIbα receptor. This interaction serves to localize αT on the platelet surface and steer the electropositive αT exosite 1 for productive interaction with the negative region of PAR1 exodomain (^53^EPFWEDEE^60^), allowing PAR1 scissile R^41^-S^42^ bond to enter the protease active site and be efficiently cleaved (60). After cleavage, the tethered positive ligand ^42^SFLLRN^47^— newly generated, couples with the negative extracellular receptor surface and triggers intracellular signalling (54). The mechanism of PAR4 proteolytic activation is very similar to that of PAR1, even though a negative patch in the receptor extracellular loop-2 region has been proposed for binding to αT exosite 2 (60).

The results reported above allow to propose that αSyn may function as a negative regulator of αT-mediated platelets activation, acting either directly, *via* competitive binding to PAR1, and indirectly, by scavenging αT on the plasma membrane. αSyn modulatory function may be accomplished mainly through charge-charge interactions (see below), acting long-range as an “electrostatic filter” to favor or disfavor binding to different platelet receptors. This should not be surprising, as αSyn contains ~30% of charged amino acid residues in its sequence. Although αSyn has a total net negative charge (pI: 4.7), the positive N-terminal and the negative C-terminal regions can interact independently, according to their specific charge state (5).

According to the model depicted in **Fig. 11**, free plasma αSyn, or αSyn released from activated platelets after PAR1 cleavage and α-granules secretion, can interact through its positive NT with the highly negative PAR1 extracellular domain (**Fig. 10B**) and prevent αT binding/activation. Concomitantly, other αSyn molecules can adhere to, *via* NT binding, and become concentrated (15,17) on the negative (phosphatidylserine rich) outer leaflet of activated platelets plasma membrane while the disordered/flexible negative C-terminal tail can favorably couple with the electropositive αT exosite 2 (**Fig. 10A**) to scavenge the protease and thus down-regulating platelets activation. Intriguingly, this scavenging effect may be particularly efficient if one considers that the affinity of αT for chip-immobilized αSyn is the medium nanomolar range (K_d_ = 44 nM) (**Fig. 5D**).

**Figure 11.**
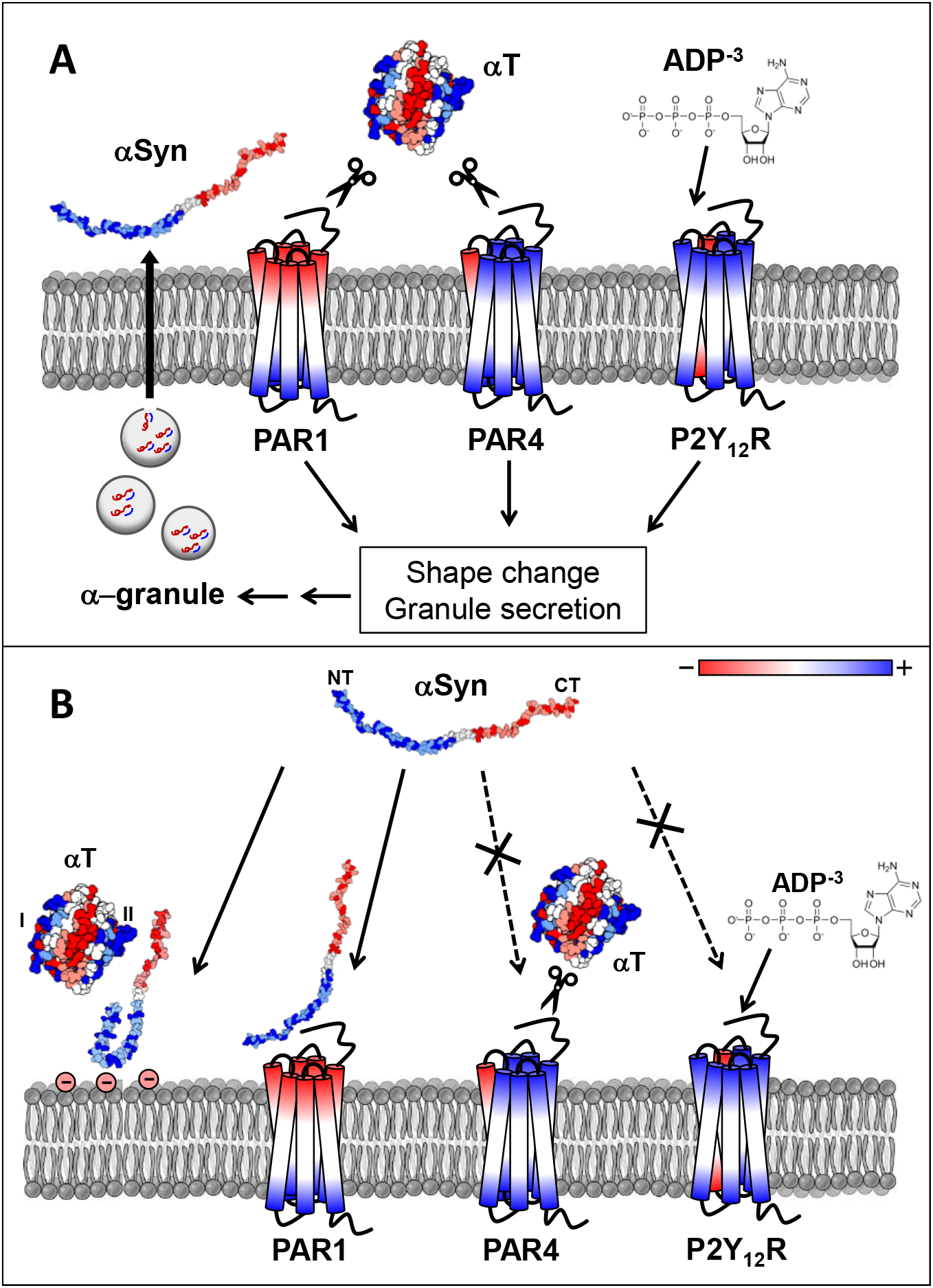
Schematic representation of the proposed mechanism of αSyn platelet antiaggregating activity. (**A**) Proteolysis of PAR1 and PAR4 exodomains by αT or direct binding of ADP to P2Y_12_R trigger intracellular signalling cascade, leading (among other effects) to platelet shape change, reversal of plasma membrane polarization, and secretion of α-granules containing αSyn. (**B**) Secreted αSyn or circulating plasma αSyn can interact through its positive NT with the highly negative PAR1 extracellular domain and impair αT binding/activation. Due to electrostatic repulsion, αSyn is not expected to interact with the positive exodomain surface of PAR4 and P2Y_12_R, allowing αT and ADP to still bind and activate the receptors. Extracellular αSyn is thought to adhere to the negative outer leaflet of activated platelets plasma membrane through its positive NT, which becomes ordered and concentrated on the membrane surface, while the negative C-terminal tail remains disordered/flexible to favorably couple with the electropositive αT exosite 2. As a result, part of αT molecules are scavenged and platelet activation is down-regulated by one of the activation products (i.e. αSyn) in a negative feedback mechanism. Blue and red colors indicate electropositive and electronegative regions, respectively, on αSyn, αT, and platelet receptors; phospholipids are in grey.

The model also explains on simple electrostatic grounds how high αSyn concentrations can completely inhibit platelet activation by TRAP6 (**Fig. 2B**), whereas at the same αSyn concentrations αT still has a significant, residual aggregating activity (~45%). Conversely to PAR1, in fact, PAR4 extracellular domain is electropositive (**Fig. 10C)** and therefore it is not expected to interact with (and be inhibited by) the positive αSyn N-terminal region. Hence, the residual platelets activating function of αT, measured in the presence of αSyn, can be reasonably assigned to the proteolytic activation of PAR4 by free αT molecules. Similar considerations also hold for explaining the poor antiaggregating effect of αSyn on ADP-induced activation (**Fig. 2D**). P2Y_12_R extracellular surface is indeed mainly electropositive, with only some interspersed negative spots, resulting in charge-charge repulsion with the positive αSyn NT. Furthermore, some sequestration of the negative ADP^−3^ agonist by the electropositive NT in the full-length αSyn cannot be ruled out (see below).

The “electrostatic model” highlighted above also provides reasonable explanation for the different platelets antiaggregating functions of αSyn(103-140) and 6xHis-αSyn(1-96) in different platelets stimulation assays (**Fig. 2B-D**). When isolated, the negative αSyn(103-140) does not affect platelets activation by TRAP6, αT or ADP^−3^, likely because of electrostatic repulsion with the negative PAR1 exodomain surface and poor stereochemical fit with PAR4 and P2Y_12_R (see Results). Likewise, binding of αSyn(103-140) to αT is driven by the electrostatic complementarity with the positive exosite 2 of the protease, while the bound peptide does not impair αT to cleave and activate PAR1 and PAR4. Noteworthy, 6xHis-αSyn(1-96) has a variable (even opposite) effect on the platelets activation assays explored in this work. In the TRAP6 activation test, 6xHis-αSyn(1-96) faithfully reproduces PAR1 inhibitory properties of intact αSyn, thus strongly supporting our hypothesis that NT may directly bind to and competitively inhibit PAR1 activation induced by TRAP6. When tested in the αT-activation assay, αSyn(1-96) displays a reduced antiaggregating activity, compared to full-length αSyn, consistent with the lack of the C-terminal tail 103-140 which, in the membrane-bound αSyn, is responsible of αT scavenging. Rather surprisingly, the antiaggregating effect of αSyn(1-96) is much increased in the ADP-activation test. This is likely caused by αSyn(1-96) binding of ADP^−3^ agonist, reducing the concentration of the agonist available for activating P2Y_12_R. This effect is much more pronounced with αSyn(1-96) than with full-length αSyn, as the former has a much higher positive charge density (pI: 9.4) compared to intact αSyn (pI: 4.7), where the presence of the negative C-terminal tail (pI: 3.1) could interfere with ADP binding. This interpretation is consistent with the known affinity of small/disordered electropositive nucleoproteins (e.g. histones and protamines) for (poly)nucleotides (61) and with very recent data showing that 12 μM protamine (51 amino acids, pI: 12), antagonises ADP activation of platelets (62). In the latter case, however, no mechanistic insight has been provided.

### Blood aSyn in health and disease

Although several different activities have been reported for αSyn (e.g. modulation of exocytosis/endocytosis, histone and metal binding, and cellular ferrireductase activity), the normal physiological function of this protein still remain to be fully elucidated (16). αSyn has been identified in all cellular components of the haematopoietic system (20,21) and in the blood (19,22-24) and recent findings, obtained with αSyn-/-knockout mice, indicate that the protein plays an important role in normal function of these cells (25). Interestingly, platelets are the main hosts of αSyn in the blood, containing 264±36 ng of αSyn *per* mg of total protein compared to 131±23 ng of the erythrocytes (23,26), while αSyn levels increase during differentiation of megakaryocytes to platelets. Altogether these findings have widened our understanding of αSyn function outside the CNS and, noteworthy, add more weight to our proposal that haematic αSyn can act as a modulator of platelet activation *in vivo* by interfering with the αT-PAR1 functional axis. Intriguingly, platelets also express high levels of other amyloid-related proteins, such as the amyloid precursor protein (APP) and Aβ(1–42) fragment (63), and β2-microglobulin (β2M) (64), which are heavily involved in the onset of Alzheimer’s disease (i.e. APP and Aβ) and neurocognitive decline (i.e. β2M). Noteworthy, elevated plasma levels of either Aβ or β2M are positively related to higher incidence of thrombosis (63,64).

The platelet antiaggregating function, herein proposed for αSyn, is keeping with earlier *in vitro* results suggesting that the protein is related to blood anticoagulant pathways *in vivo*. For instance, exogenous αSyn has been shown to pass across the plasma membrane and impair calcium-dependent release of α-granules from platelets (27), and to act as a negative regulator of the exocytosis of von Willebrand factor (vWF) from vascular endothelial cells (65), where vWF recruits and activates platelets in primary haemostasis (28). Importantly, the lack of platelet αSyn, as in α-syn-/-knockout mice, results in a hypercoagulable phenotype (25,66).

Our results are also consistent with *in vivo* procoagulant and anticoagulant clinical settings where αSyn levels are either decreased (i.e. aging) or increased (i.e. PD) and for which an opposite trend of thrombotic events have been observed. Although it is difficult to establish a direct causative link between the concentration of a given (protein) molecule and the onset of a complex multifactorial physio-pathological state, blood αSyn levels are markedly decreased in elderly people (67) and aging is widely recognized as a primary risk factor for thrombosis (68). On the other hand, very recent data indicate that αSyn plasma concentration (but not platelet or erythrocyte αSyn levels) is >20-fold higher in PD patients than in healthy individuals (69) and, as a matter of fact, earlier studies seem to indicate that ischemic stroke, myocardial infarction or coronary arterial disease are significantly less frequent in PD patients than in healthy controls (70–72) while platelets from PD patients are less sensitive to aggregating agents like αT and ADP (73). Even though epidemiologic and neurochemical facets of Parkinson’s disease might allow to envisage some benefit or protection against thrombotic events, other studies suggest that the prevalence of thrombotic disorders is higher in PD patients than in controls (74,75). The results of these studies, however, are likely affected by a confounder bias, as the enrolled PD patients had a history of anti-Parkinson medication with Levodopa, known to increase plasma levels of homocysteine which is recognized as thrombotic risk factor (76). Noteworthy, in a retrospective case-control study, newly diagnosed PD patients at early disease stage (not treated with Levodopa) were characterized by a significantly lower prevalence of vascular risk factors than matched controls (77).

In conclusion, the results reported in this study put forward a novel physiological function of αSyn, which unfolds outside the CNS, to downregulate αT-induced platelet activation. Further studies are needed to address the impact of αSyn antiaggregating function, in comparison with that of other platelet-derived amyloidogenic proteins (i.e. APP/Aβ and β2M), at the interplay of amyloidosis and thrombosis.

### Experimental procedures

#### Reagents

Human plasma αT (EC 3.4.21.5) and ProT were purchased from Haematologic Technologies (Essex Junction, VT, USA). Ecarin from *Echis carinatus* venom, bovine pancreas trypsin, human plasma fibrinogen, Ac-Tyr-NH_2_, Ac-Phe-NH_2_, fluorescein isothiocyanate and PABA were purchased from Sigma (St. Louis, MO, USA). The chromogenic substrate S2238 was from Chromogenix (Milan, Italy). Hirugen (^54^GDFEEIPEEY*LQ^65^) and [F]-hirugen (46), fibrinogen γ’–peptide (^408^VRPEHPAETEY*DSLY*PEDDL^427^) (78), PAR1(38–60) (^38^LDPR↓SFLLRNPNDKYEPFWEDDE^60^) (45), Hir(1–47) (79), and αSyn(103-140) were synthesised by standard N^α^-fluorenylmethyloxycarbonyl solid-phase chemistry on a PS3 automated synthesizer (Protein Technologies Int., Tucson, AZ, USA), purified to homogeneity (>98%) by semipreparative RP-HPLC, and characterized by high-resolution mass spectrometry. Notably, Y* indicate phosphorylated Tyr-residues. N^α^-Fmoc-protected amino acids, solvents and reagents for peptide synthesis were purchased from Applied Biosystems (Forster City, CA, USA) or Bachem AG (Bubendorf, Switzerland). Salts, solvents and other reagents were of analytical grade and purchased from Sigma or Fluka (Darmstadt, Germany).

#### Production and characterization of recombinant αSyn species

All recombinant human synuclein derivatives (i.e. αSyn, 6xHis-αSyn, 6xHis-αSyn(1-96), and αSyn-GFP), were produced as previously detailed (31,32). Briefly, BL21*(DE3) pLysS *Escherichia coli* cells were transformed, using the heat-shock method, with *p*RSET-B plasmid containing human αSyn gene and selected on a Luria-Bertani (LB) Agar Amp^+^ (0.1 mg/ml) solid culture medium overnight. Transformed cells were grown at 37°C in LB Broth Amp^+^ (0.05 mg/ml) and induced (O.D. = 0.6) with isopropyl β-D-1-thiogalactopyranoside (IPTG, 0.1 mg/ml) under vigorous shaking. For αSyn and αSyn-GFP, after 3-h induction with IPTG, bacteria were harvested by centrifugation (6.000 rpm, 15 min, at 4°C), and the pellet sonicated in 40 mM Tris-HCl, pH 8.0, 0.1 M NaCl (buffer A). After 10-min boiling, the suspension was centrifuged (12.000 rpm, 10 min, 4°C). The supernatant, containing soluble αSyn or αSyn-GFP, was dialyzed overnight at 4°C against buffer A, containing 2 mM EDTA. For 6xHis-αSyn and 6xHis-αSyn(1-96), after sonication in buffer A, the recombinant proteins was purified by IMAC. The bacterial lysis supernatant (50 ml) was loaded onto a fast-flow Ni^2+^-IMAC (1 x 3 cm) HiTrap column, using a model P-1 peristaltic pump (Pharmacia, Uppsala, Sweden) at a flow-rate of 0.1 ml/min. The flow-through was discarded and the column connected to an Äkta-purifier system (Marlborough, MA, USA). After washing with buffer A (60 ml), 6xHis-tagged proteins were eluted from the column (0.5 ml/min) with buffer A, pH 6.5, containing 0.4 M imidazole. The material eluted in correspondence of the major chromatographic peak was collected and dialyzed overnight at 4°C against phosphate buffered saline, pH 7.4. Recombinant proteins were further purified by RP-HPLC on a C18 semi-preparative column (10 x 250 mm, 5μm, 300Å) from Grace-Vydac (Hesperia, CA, USA), eluted with a linear acetonitrile-0.078% trifluoroacetic acid gradient at a flow rate of 1.5 ml/min. After lyophilisation, recombinant proteins in water:acetonitrile (1:1 v/v), containing 1% formic acid were characterized by high-resolution mass spectrometry using a Waters (Milford, MA, USA) Xevo-G2S Q-TOF spectrometer. To obtain purified αSyn in the monomeric state, the lyophilized protein (1 mg) was dissolved in 2 mM NaOH (100 μl) and 1 M NaOH (10 μl), up to pH 11.0. After centrifugation (15.000 rpm, 15 min), the supernatant was withdrawn and added with 0.1 M Tris-HCl, pH 7.0 (200 μl), down to pH 8.0. Freshly dissolved αSyn samples were used for further spectroscopic and functional analyses. Purified αSyn solutions were divided into aliquots, lyophilized, and stored at −20°C. After thawing in ice-water bath, αSyn aliquots were immediately used for subsequent functional/binding analyses.

#### Production and characterization of thrombin derivatives

The plasmid containing the cDNA of prethrombin-2 was a generous gift of Prof. James A. Huntington (University of Cambridge, Cambridge, UK). The recombinant inactive mutant S195A, obtained by single-point mutagenesis, was expressed in *E. coli*, subjected to *in vitro* disulphide oxidative folding, activation by ecarin, and characterized as previously detailed (45,48). β_T_-thrombin (β_T_T) was obtained by proteolysis of human αT (7 μM) with bovine pancreas trypsin (35 nM) for 3 hours at 37°C in HBS, i.e. 4-(2-hydroxyethyl)-1-piperazineethanesulfonic acid (HEPES) buffered saline, pH 7.4, and characterized as previously detailed (41,46).

#### Platelets aggregation assays

The effect of αSyn species on platelets aggregation induced by αT, TRAP or ADP was measured at 37°C in whole blood and PRP by MEA, using a multiplate analyzer (Dynabyte, Munich, Germany) (36,41). Citrate-treated venous blood samples were taken from five healthy donors: two males and three females, 28–35 years of age, and non-smokers. The donors gave written informed consent for participation in this study, which was approved by the Institutional Ethics Committee of the Padua University Hospital. PRP was obtained from three healthy subjects as previously described (80,81). Briefly, after centrifugation of citrated venous blood samples (250 g, 10 min, 4°C), the supernatant was diluted 5:1 (v/v) with PBS, pH 7.4, containing 10 mM EDTA, and centrifuged (12.000 g, 1 min, 4°C) to allow sedimentation of platelets. The pellet was washed twice with PBS/EDTA and finally resuspended in HBS, pH 7.4. Platelet count were determined with a Cell-Dyn Emerald 22 cytometer from Abbott Diagnostics (Chicago, IL, USA). Increasing concentrations (0-20 μM; 300 μl in HBS) of monomeric samples of αSyn species, i.e. full-length αSyn, 6xHis-αSyn(1-96) and αSyn(103-140), were pre-incubated (30 min, 37°C) with whole blood or PRP (300 μl, 160.000–200.000 platelets/μl). Platelets aggregation was started by addition of TRAP6 or ADP stock solutions (20 μl) and analysed by MEA over 10-min reaction time. ADP-test and TRAP-test solutions for Multiplate assays were purchased from Roche Diagnostics (Basel, Switzerland). When the effect of αSyn species on αT-induced aggregation was measured, protease solutions (20 μl in HBS) were pre-incubated (30 min, 37°C) with increasing αSyn concentrations (0-20 μM, 300 μl in HBS) and then added to blood or PRP samples (300μl). For each MEA measurement, the area under the aggregation curve (AUC) was determined for single donors and the average value expressed as %AUC, relative to the value determined in the absence of αSyn (AUC_0_) (45,48).

#### Fibrin generation assays

Fibrin generation was started by adding αT (1 nM) to a freshly desalted Fb solution (0.44 μM) in HBS at 37°C, while the time course of clot formation was followed by continuously recording the solution absorbance at 350 nm (i.e. the turbidity) on a double-beam V-630 Jasco (Tokyo, Japan) spectrophotometer (41,45,48). The effect of αSyn was estimated by first incubating αT with increasing concentrations (0-20 μM) of αSyn and then adding a desalted Fb solution.

#### Enzymatic activity assays

Hydrolytic activity of αT was determined at 37°C in HBS on the chromogenic substrate S2238 by measuring the release of pNA at 405nm (ε^M^_405nm_= 9920 M^−1^·cm^−1^). The kinetics of FpA and FpB release was followed as earlier reported (41,48). Briefly, human fibrinogen (Fb) (ε^M^_280nm_ = 5.1·10^5^ M^−1^·cm^−1^) was desalted on an in-house packed: (8×125mm) G10 fast-flow column (GE Healthcare, Chicago, IL, USA) eluted with HBS, pH 7.4, at a flow-rate of 0.3 ml/min. Freshly prepared Fb (0.35 μM) was reacted at 37°C with human αT (300 pM) in the presence of 15 μM αSyn and at fixed time points proteolysis mixtures were added with formic acid (2% v/v final concentration) to block the proteolysis reaction and induce precipitation of unreacted Fb. After centrifugation (10.000 g, 5 min, 4°C), the supernatant (1.0 ml) was withdrawn, lyophilized, dissolved in 6 M guanidinium hydrochloride solution (170 μl) and injected (100 μl) onto a RP-HPLC (4.6 x 250mm) C18 column (Grace-Vydac, Columbia, MD, USA). The column was equilibrated with 40 mM ammonium phosphate buffer, pH 3.1, and eluted with an acetonitrile gradient. The absorbance of the effluent was recorded at 205 nm and the amount of FpA (ε^M^_205nm_ = 4.40·10^4^ M^−1^·cm^−1^) and FpB (ε_MM_= 5.12·10^4^ M^−1^·cm^−1^) released was determined by integrating the area under the chromatographic peaks. A LC-4000 HPLC system (jasco, Tokyo, Japan) was used for all analyses.

The specificity constants, k_cat_/K_m_, for the release of fibrinopeptides were determined by interpolating the data points to equations 1 and 2 (41,48):

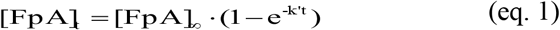

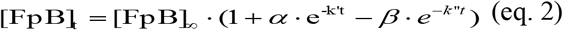

where [FpA]_t_ or [FpB]_t_ and [FpA]_∞_ or [FpB]_∞_ are the concentration of FpA or FpB at time t and ∞, respectively, and k′ and k′′ are the observed kinetic constants for FpA or FpB release, obtained as fitting parameters. Under pseudo-first order conditions and low substrate concentration, the specificity constants could be easily determined k_catA_/K_mA_ = k′/[E] and k_catB_/K_mB_ = k′′/[E], where [E] is the protease concentration.

Hydrolysis of the synthetic peptide PAR1(38–60) (1 µM) by αT (150 pM) was carried out at 25°C in TBS, in the presence of αSyn (15 μM). At time points, aliquots (360 μl) were taken, acid quenched (10 μl, 4% aqueous TFA) and loaded (350 µL) onto 99.360 a Grace-Vydac (4.6 x 250 mm) C18 column. The column was eluted with a linear acetonitrile-0.078% TFA gradient from 10-45% in 40 min and the release of PAR1(42–60) (ε^M^_205nm_ = 95870 M^−1^·cm^−1^) was quantified by integrating the area under the chromatographic peak. The kinetic data were interpolated with equation 3, describing a pseudo-first order reaction (41,46,48)

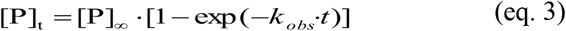

where [P]_∞_ is the concentration of the fragment PAR1(42–60) when the proteolysis reaction was complete and k_obs_ is the observed kinetic constant for PAR1(38–60) hydrolysis, obtained as a fitting parameter. As for the release of fibrinopeptides, under pseudo-first order conditions and low substrate concentration, k_cat_/K_m_ could be derived as k_obs_/[E].

#### Spectroscopic methods

*Ultraviolet absorption spectroscopy*. The concentrations of protein/peptide solutions were determined by measuring the absorbance at 280 nm on a Jasco V-630 double-beam spectrophotometer, using the following molar absorptivity values (ε^280^): plasma αT and β T, 67.161 M^−1^·cm^−1^; recombinant rS195A, 66.424 M^−1^cm^−1^; ProT, 99.360 M^−1^·cm^−1^) M^−1^·cm^−1^; αSyn, 5.960 M^−1^·cm^−1^; 6xHis-αSyn(1-96), 1.490 M^−1^·cm^−1^; αSyn(103-140), 4.470 M^−1^·cm^−1^; hirugen, 418 M^−1^·cm^−1^; Hir(1–47), 3.355 M^−1^·cm^−1^; fibrinogen γ’–peptide, 837 M^−1^·cm^−1^; [F]-hirugen at 492 nm, 68.000 M^−1^·cm^−1^; PABA at 336 nm, 548 M^−1^·cm^−1^; S2238 at 316 nm, 12.700 M^−1^·cm^−1^. The concentration of active αT was also determined by active-site titration with hirudin (50) and found identical (±5%) to that determined spectrophotometrically.

#### Dynamic light scattering

Measurements were performed at 37°C on a Zetasizer-Nano-S instrument (Malvern Instruments, Worchestershire, UK) at a fixed angle (i.e. 173°) from the incident light (i.e. He–Ne 4 mW laser source at 633 nm). Polystyrene cuvettes (1-cm pathlength, 100 μl) (Hellma, Switzerland) were used for all measurements. Each measurement consisted of a single run (15 s). Scattering data were analyzed with the Nano-6.20 software and expressed as percentage of volume size distribution, from which the value of *d*_*H*_ and %PD were extracted (46,82), where *d*_*H*_ is the diameter of a hard sphere that diffuses at the same speed as the molecule being measured, and %PD is the width of the particle size distribution of a protein in a given sample.

#### Fluorescence spectroscopy

*B*inding measurements were carried out at 37°C in HBS, containing 0.1% PEG-8000 (w/v), on a Jasco FP-6500 spectrofluorimeter. Aliquots (2-10 μl) of αSyn or αSyn(103-140) in HBS were added, under gentle magnetic stirring, to an αT solution (70 nM) in the same buffer. At each ligand concentration, samples were incubated for 2 min at 37°C and excited at 295 nm, using an excitation/emission slit of 5/10 nm. Fluorescence intensity was recorded at 334 nm, i.e. the λ_max_ of αT emission, after subtracting the corresponding spectra of the ligands alone. Fluorescence data were corrected for sample dilution (<5%). To prevent photobleaching of Trp-residues, a 1-cm pathlength quartz cuvette (2 ml) with two frosted walls, diffusing the incident light inside the sample, was used. The optical density of the solution was always kept <0.05 units both at λ_ex_ and λ_em_, to avoid inner filter effect (47). A similar procedure was used for measuring the affinity of all other site-specific ligands tested in this work (i.e. PABA, S2238, Hir(1–47), hirugen, [F]-hirugen and fibrinogen γ’–peptide) for αT in the presence of constant, saturating αSyn or αSyn(103-140) concentration (20 μM). When the binding of PABA was being studied, samples were excited at 336 nm and the emission of PABA was recorded at 375 nm, after baseline subtraction and correction for inner filter effect, as detailed elsewhere (47). For [F]-hirugen binding, aliquots of αT S195A mutant stock solution (30 μM) were incrementally added to a [F]-hirugen solution (60 nM). Samples were excited at 492 nm and the decrease of fluorescence intensity of [F]-hirugen was recorded at 516 nm as a function of αT (46).

The data points were interpolated with equation 4, describing the single-site binding model R + L ↔ RL (47):

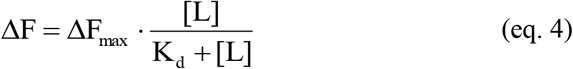

where L is the ligand concentration, ΔF and ΔF_max_ are the changes of fluorescence intensity measured at intermediate or saturating ligand concentrations, while the dissociation constant, K_d_, was obtained as a fitting parameter. For Hir(1–47) and [F]-hirugen binding to αT, fluorescence data were interpolated with equation 5, describing the tight-binding model (47).

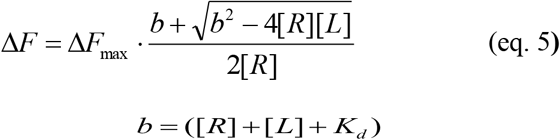

where [R] and [L] are the total enzyme or ligand concentrations.

#### Surface plasmon resonance

SPR analyses were carried out on a dual flow-cell Biacore X-100 instrument from GE Healthcare. 6xHis-αSyn was noncovalently immobilized onto a Ni^2+^-chelated nitrilotriacetate (NTA) carboxymethyldestrane sensor chip and incremental concentrations of S195A were loaded. The Ni^2+^-NTA/6xHis-αSyn chip assembly was prepared as follows: the NTA chip (GE Healthcare) was first washed (flow-rate: 30 μl/min) with 0.35 M EDTA, pH 8.3 (contact time: 700 sec) and then loaded with 0.5 mM NiCl_2_ solution (contact time: 400 sec); excess Ni^2+^ was removed by injecting 3 mM EDTA solution (contact time: 350 sec), whereas non-chelating NTA-groups were irreversibly blocked with ethanolamine, after carboxylate activation (contact time: 800 sec) with N-(3-dimethylaminopropyl)-N′-ethylcarbodiimide and N-hydroxysuccinimide; finally, a solution of 6xHis-αSyn (200 nM) was injected on the sensor chip (contact time: 400 sec) to yield a final immobilization level of 2194 response units (RU). The Ni^2+^-NTA/6xHis-αSyn sensor chip was challenged (flow-rate: 30 μl/min; contact time: 350 sec) with increasing concentrations of inactive S195A thrombin mutant, β_T_T, and ProT. All measurements were carried out at 37°C in HBS-EP^+^ buffer (10 mM HEPES, pH 7.4, 0.15 M NaCl, 50 μM EDTA, 0.005% v/v polyoxyethylene sorbitan). Between two consecutive runs, the regeneration of Ni^2+^-NTA/6xHis-αSyn chip was achieved with HBS-EP^+^ buffer, containing 2 M NaCl. Each sensogram was subtracted for the corresponding baseline, obtained on the reference flow cell and accounting for nonspecific binding, i.e. typically less than 2% of RU_max_. The binding data were analyzed using the BIAevaluation software. The dissociation constant (K_d_) relative to the binding of αT to immobilized αSyn was obtained as a fitting parameter by plotting the RU value at the steady state (RU_eq_) *versus* [αT] and interpolating the data points with equation 6, describing 1:1 binding model:

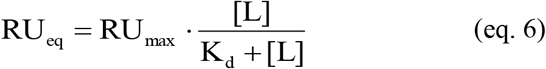

where L is the concentration of αT, while RU_eq_ and RU_max_ are the RU values measured (at the steady state) with intermediate or saturating [L] (41,48).

### *Isothermal titration calorimetry* (ITC)

ITC titrations were performed at 25±0.1°C in 20 mM HEPES pH 7.4, 0.15M, using a MicroCal VP-ITC instrument, as described (83). To a S195A thrombin mutant solution (1.7 ml, 2 µM) were sequentially added 25 aliquots (10 μl each) of αSyn stock solution (40 μM), under continuous stirring (307 r.p.m.) and a delay of 4 min after each injection. Before analysis, protein samples were dialyzed overnight in the same buffer, using a Slide-A-Lyzer (3.5-kDa cutoff) from ThermoFischer Scientific (Waltham, MA, USA), and thoroughly degassed. The heat of dilution was determined in control experiments by injecting aliquots (10 µl) of αSyn stock solution (40 µM) into buffer and this was subtracted from the integrated binding isotherm prior to curve fitting. The thermograms were analysed using the MicroCal ITC Data Analysis software.

### Fluorescence microscopy techniques

PRP was prepared as described above (80,81). Afterwards, isolated platelets in serum-free Iscove’s Modified Dulbecco’s medium were seeded (20×10^6^ platelets/well) in 24-well culture plates containing a glass coverslip coated with gelatine. After 24 h, resting platelets were incubated with increasing concentrations of αSyn-GFP (0-2 μg) in HBS. In the same experiment, resting platelets were first activated with 10 µM TRAP6 and then added with αSyn-GFP. Untreated resting platelets and TRAP6 stimulated platelets, without αSyn-GFP, were used as controls. Resting and stimulated platelets were observed for 30 min at 37°C, under 5% CO_2_ flow, in the time-lapse mode using a DMI6000-CS fluorescence microscope (Leica Microsystem, Wetzlar, Germany), equipped with a DFC365FX camera and a 40x/0.60 dry objective magnification. Images were taken in real time (at 30 min) using a differential interference contrast (DIC) and fluorescence objectives and processed using the Leica Application Suite 3.1.1. software (Leica Microsystem).

In another experiment, both resting and activated platelets were incubated for 24 h with αSyn-GFP (0.7 and 1.4 μM), washed twice with PBS and fixed for 20 min in 2% paraformaldehyde. The slides were mounted with Mowiol antifade solution (Sigma-Aldrich, St. Louis, MO, USA) and directly observed with a DMI6000-CS microscope. Images were acquired using 100x/1.4 oil immersion objective magnification. Finally, endogenous αSyn exposure on platelets plasma membrane was detected by immunofluorescence microscopy, using the same equipment as above. Resting and TRAP6 activated platelets were incubated for 1 h at 37°C with 5 μg/ml mouse anti-human αSyn monoclonal antibody [α-synuclein(211):sc-12767] form Santa Cruz Biotechnology (Dallas, TX, USA), followed by addition of a diluted (1:200) Alexa Fluor 594-conjugated goat anti-mouse IgG (Thermo-Fisher, Waltham, MA, USA). Both primary and secondary antibodies were diluted in PBS, containing 0.5% bovine serum albumin. Unspecific binding was assessed by incubating platelets with the secondary antibody alone, in the absence of the primary antibody.

### Computational methods

PAR4 (UniProt code: Q96RI0; amino acid residues Asp65-Phe347) structure was modeled by homology on the template structure of PAR1 (PDB code: 3vw7; UniProt code: P25116; amino acid residues Asp91-Cys378) (54) with which it shares 34.6% sequence identity and 56.1% sequence similarity. The Swiss-Model software was used (84). Electrostatic potential calculations were performed using APBS software (53). For αT, calculations were run on the non-glycosylated X-ray structure of αT (1ppb), after removal of the coordinates of the inhibitor D-Phe-Pip-Arg-chloromethylketone, water and HEPES molecules (49). The coordinates of human PAR1 (3vw7) (54) and P2Y_12_ receptor (4ntj) (55) bound to the inhibitors Vorapaxar and AZD1283, respectively, were considered. The electrostatic contribution of Na^+^-ion bound to PAR1 was not considered in our calculations. A solvent dielectric of 78.14 and a protein dielectric of 2.0 at 310K in 150 mM NaCl were used. Final electrostatic maps were constructed by subtracting the protein self-energies from the calculated map using the DXMATH utility in APBS. Notably, to facilitate crystallogenesis, T4 lysozyme (T4L) and the BRIL domain were inserted into the intracellular loop 3 of PAR1 (54) and P2Y_12_R (55), respectively. In the recombinant PAR1-T4L fusion protein the N-terminal exodomain was missing. The coordinates of the bound inhibitor were virtually removed, along with the inserted structure of T4L and BRIL. To minimize artefactual charge perturbations, following virtual domain excision, the remaining N- and C-termini were made neutral by acetylation or amidation.

## Data availability

All other data that support the findings of this study are available from the corresponding author upon reasonable request.

## Author contributions

G.P., L.A., I.A., A.P. and F.U. performed research; C.M.R. and P.S. performed fluorescence microscopy work; D.P. performed modelling work and electrostatic calculations; A.N. produced recombinant αSyn species; V.D.F. inspired and coordinated the work, analyzed and interpreted the data, and wrote the manuscript; all authors analyzed and interpreted the data and reviewed the final content of the manuscript.

## Acknowledgements

Part of this work was presented at the ISTH-2017 conference, July 8-13, 2017 - Berlin (Germany). Commun. PB-1706. The authors are grateful to Dr. Daniele Dalzoppo (University of Padua) for critically reading the manuscript and Dr. Nicola Pozzi for performing some very preliminary measurements. The authors also thank Dr. Vittorio Pengo (University of Padua) for providing accessibility to the Multiplate analyser at the beginning of this study and for supplying us with some blood samples from normal subjects. The generous gift of the plasmid containing the cDNA of prethrombin-2 by Prof. James A. Huntington (University of Cambridge, Cambridge, UK) is gratefully acknowledged.

## Abbreviations

αSyn: recombinant human α-synuclein
αSyn(1-96): recombinant polypeptide corresponding to αSyn sequence 1-96
αSyn(103-140): synthetic peptide corresponding to αSyn sequence 103-140
αSyn-GFP: αSyn fused at the C-terminal end with GFP
αT: human α-thrombin
ADP: adenosine 5’-diphosphate
AUC: area under the aggregation curve
DLS: dynamic light scattering
EDTA: ethylenediaminetetraacetate
Fb: human fibrinogen
GFP: green fluorescent protein
HBS 20 mM HEPES pH 7.4: 0.15 M NaCl, 0.1 % PEG-8000 (w/v)
HBS-EP^+^: 10 mM HEPES
pH 7.4, 0.15 M NaCl: 50 μM EDTA, 0.005% (v/v) polyoxyethylene sorbitan
HEPES: 4-(2-hydroxyethyl)-1-piperazineethanesulfonic acid
IMAC: immobilized metal ion affinity chromatography
ITC: isothermal titration calorimetry
MEA: Multiple Electrode Aggregometry
NTA: nitrilotriacetate
PABA: *p*-aminobenzamidine
PAR1: protease-activated receptor-1
PBS: phosphate buffered saline
*p*NA: *p*-nitroaniline
PD: Parkinson’s disease
ProT: human prothrombin
PRP: platelet rich plasma; S2238, (D)-Phe-Pip-Arg-*p*-nitroanilide
SPR: surface plasmon resonance
TBS: 5 mM Tris-HCl pH 7.4, 0.15 M NaCl
TFA: trifluoroacetic acid
TRAP6: thrombin receptor activating peptide, having the sequence SFLLRN-NH_2_
Tris-HCl: Tris(hydroxymethyl)aminomethane hydrochloride
vWF: von Willebrand factor

